# Complementary Ribo-seq approaches map the translatome and provide a small protein census in the foodborne pathogen *Campylobacter jejuni*

**DOI:** 10.1101/2022.11.09.515450

**Authors:** Kathrin Froschauer, Sarah L. Svensson, Rick Gelhausen, Elisabetta Fiore, Philipp Kible, Alicia Klaude, Martin Kucklick, Stephan Fuchs, Florian Eggenhofer, Susanne Engelmann, Rolf Backofen, Cynthia M. Sharma

## Abstract

**Background:** Knowing the molecules encoded by bacterial pathogens and how their expression is regulated is essential to understand how they survive, colonize, and cause disease. RNA-seq technologies that map transcriptomes have revealed a wealth of new transcripts in bacterial pathogens. However, they do not provide direct evidence or coordinates for coding potential. In particular, they miss small proteins (≤ 50-100 amino acids) translated from small open reading frames (sORFs). However, this still poorly annotated component of bacterial genomes shows emerging roles in bacterial physiology and virulence.

**Results:** Here, we present an integrated approach based on complementary ribosome profiling (Ribo-seq) techniques to map the “translatome” of *Campylobacter jejuni*, the most common cause of bacterial gastroenteritis. Besides conventional Ribo-seq, we employed translation initiation site (TIS) profiling to map start codons and reveal internal sORFs. We also developed a Ribo-seq approach for mapping of translation termination sites (TTS), which revealed stop codons not apparent from the reference genome in virulence-associated loci. Our translatome map confirms translation of leaderless ORFs and leader peptides, re-annotates start or stop codons of 35 genes, and reveals isoforms generated by internal start sites. It also adds 42 novel sORFs in diverse contexts to the *C. jejuni* annotation, such as within small RNAs, in 5′ untranslated regions (UTRs), or internal/out-of-frame/antisense in larger ORFs. Using epitope tagging/western blot and mass spectrometry we validated expression of almost 60 annotated and novel sORFs, including *cioY*, which we show encodes a conserved, 34 amino acid component of the CioAB terminal oxidase.

**Conclusions:** Overall, we provide a blueprint for integrating several Ribo-seq approaches to refine and enrich bacterial annotations.

## BACKGROUND

A complete census of the coding and non-coding features of bacterial genomes is essential to understand how bacteria survive and adapt to environmental challenges. Especially for pathogens this is important to understand how they colonize and cause disease in order to develop new antimicrobial strategies. Next generation DNA and RNA-sequencing (RNA-seq) technologies that map gene function and expression genome-wide have rapidly expanded our catalog of factors that might influence infection (Barquist and Vogel, 2015). This includes hundreds of potential small non-coding regulatory RNAs (sRNAs) in intergenic regions, which act as key regulators of bacterial stress responses, fitness, antibiotic resistance, and virulence (Felden and Augagneur, 2021; Hör et al., 2020; Svensson and Sharma, 2016; Westermann, 2018). More recently, the concept of genes within genes has emerged in different forms, *e*.*g*., as sRNAs derived from mRNA untranslated regions (Ponath et al., 2022), dual-function (coding and regulatory) sRNAs (Raina et al., 2018), or alternative open reading frames (ORFs) generated by internal start codons (Adams and Storz, 2020; Meydan et al., 2018; Orr et al., 2020). Therefore, bacterial genomes likely still hide genes with key roles in physiology and/or virulence.

A major gap in our catalog of potential factors affecting bacterial physiology and virulence is so-called ‘small proteins’, representing the small proteome. Several apparently ‘non-coding’ novel transcripts discovered by RNA-seq appear to encode short ORFs (sORFs) and are rather small mRNAs. Small proteins (here defined as ribosomally-synthesized polypeptides ≤70 amino acids (aa)), have long been overlooked or even discarded from bacterial genome annotations. However, they are emerging as important players in diverse processes in bacterial cells, including stress response, virulence, and metabolism, as well as in the biology of their phages (Ardern, 2021; Duval and Cossart, 2017; Gray et al., 2022). Many of the few functionally characterized small proteins bind and modulate the activity of regulators, enzymes, and transport complexes (Hemm et al., 2020; Orr et al., 2020; Steinberg and Koch, 2021). Recent studies suggest that even in genomes of well-studied bacteria, such as *E. coli* and the model pathogen *Salmonella*, small genes with key functions are still hiding (Venturini et al., 2020; Weaver et al., 2019), suggesting that many sORFs are yet to be discovered in diverse prokaryotes.

Searches for sORFs using comparative genomics across bacterial species and even their phages have revealed thousands of potential candidates (Durrant and Bhatt, 2021; Fremin et al., 2022; Miravet-Verde et al., 2019; Sberro et al., 2019). Validating these predictions, however, is challenging as small proteins can be difficult to detect by mass spectrometry (MS) approaches despite the development of new methodologies tailored for small proteins (reviewed in (Ahrens et al., 2022)). Ribosome profiling (Ribo-seq) harnesses the sensitivity and resolution of RNA-seq to assay the ‘translatome’ as a proxy for protein expression (Ingolia et al., 2009). Ribo-seq is based on deep sequencing of the ∼30 nucleotide(nt)-long mRNA ‘footprints’ that are protected from nuclease digestion by translating ribosomes, which identifies and measures translated transcripts and maps ORF boundaries genome-wide. While initially applied to study translational regulation, Ribo-seq unexpectedly revealed potentially widespread translation of sORFs in all kingdoms of cellular life and even viruses (Brar and Weissman, 2015; Ingolia et al., 2019), and is also a powerful method to mine the small proteome of bacterial species (reviewed in (Vazquez-Laslop et al., 2022)).

Due to technical limitations, bacterial Ribo-seq data often lack a 3-nucleotide periodicity observed for eukaryotes (Mohammad et al., 2019), which can hamper precise annotation of ORF boundaries. However, recently Ribo-seq variations that enrich footprints from initiating ribosomes have been developed to map translation initiation sites (TIS) and facilitate discovery of new ORFs and genes within genes in bacteria (reviewed in (Vazquez-Laslop et al., 2022)). TIS profiling can reveal alternative ORFs such as isoforms generated by internal in-frame TIS or ORFs nested out-of-frame in longer ORFs (Meydan et al., 2018). In bacteria, start codon mapping relies on antibiotics such as the pleuromutilin retapamulin (Ret) or the proline-rich antimicrobial peptide (PrAMP) oncocin 112 (Onc) to stall initiation complexes (Meydan et al., 2019; Weaver et al., 2019). Alternative ORFs can also arise from premature stop codons generated by mutation (*e*.*g*., phase-variation) or even frameshifting (Atkins et al., 2016). For example, the *E. coli* CopZ copper chaperone (70 aa) is generated by frameshifting within the longer *copA* gene (Meydan et al., 2017). Prediction of such proteins from genome sequences is not trivial as the ORF cannot be directly inferred from its start codon. Therefore, genome-wide experimental detection of translation termination sites (TTS) might reveal more examples and would provide further confidence to the annotation of novel sORFs.

*Campylobacter jejuni* is the most common cause of bacterial gastroenteritis worldwide and is also associated with several secondary neuropathies (Burnham and Hendrixson, 2018; Havelaar et al., 2015). Despite its success in causing human disease, its annotated genome lacks homologs of key virulence factors used by other enteric pathogens, and little is known about how it causes disease (Burnham and Hendrixson, 2018; Parkhill et al., 2000). *C. jejuni* strains show high genotypic and phenotypic diversity and RNA-seq has provided insights into its transcriptome structure and revealed several sRNAs (Dugar et al., 2013; Porcelli et al., 2013; Taveirne et al., 2013). For example, we previously generated primary transcriptome maps for four *C. jejuni* strains (Dugar et al., 2013), which revealed conserved and infection-relevant sRNAs, strain-specific promoter usage, and potential variation in ORF lengths (Alzheimer et al., 2020; Svensson and Sharma, 2021). In contrast, the *C. jejuni* translatome and small proteome have not yet been refined.

Here, we have employed three Ribo-seq methods to map the *C. jejuni* translatome. In addition to ‘standard’ Ribo-seq and TIS profiling, we also developed a new method leveraging the PrAMP apidaecin (Api) to stall terminating ribosomes at stop codons to map TTS. We integrated datasets from these methods to completely refine the *C. jejuni* ORF annotation, including start and stop codon positions, leaderless transcripts, and leader peptides. Extensive investigation of the annotated sORFs in our Ribo-seq data as well as independent validation by epitope tagging and MS validated translation of 47 out of 55 annotated sORFs ≤70 aa. We also expanded the *C. jejuni* small proteome by almost 2-fold (42 new sORFs) and revealed small proteins encoded in diverse genomic contexts including overlapping longer ORFs. Our data also revealed CioY (34 aa), which we show is a small protein component of the CioAB terminal oxidase. Overall, we add a new method to the bacterial translatomics toolkit, which can be applied to other diverse species, and provide a blueprint for approaching translatome refinement and small proteome discovery in diverse prokaryotes.

## RESULTS

### Ribo-seq in *C. jejuni* distinguishes between coding and non-coding regions

To identify and map translated *C. jejuni* ORFs, we established three complementary Ribo-seq approaches for *C. jejuni* strain NCTC11168 grown under standard conditions (log phase in rich media), as well as performed RNA-seq to measure the transcriptome in parallel (**Fig. 1A**). In addition to canonical Ribo-seq where we stalled ribosomes using either the elongation inhibitor chloramphenicol (Cm) and/or rapid chilling, we applied TIS profiling using Ret (“Ribo-RET”) and Onc treatment prior to polysome isolation to map start codons (Meydan et al., 2019; Weaver et al., 2019). Furthermore, we developed a new approach to detect TTS (stop codons) genome-wide using apidaecin-137 (Api). The analysis and re-annotation of the *C. jejuni* coding genome using our datasets is described in the subsequent sections of this study.

**Figure 1.**
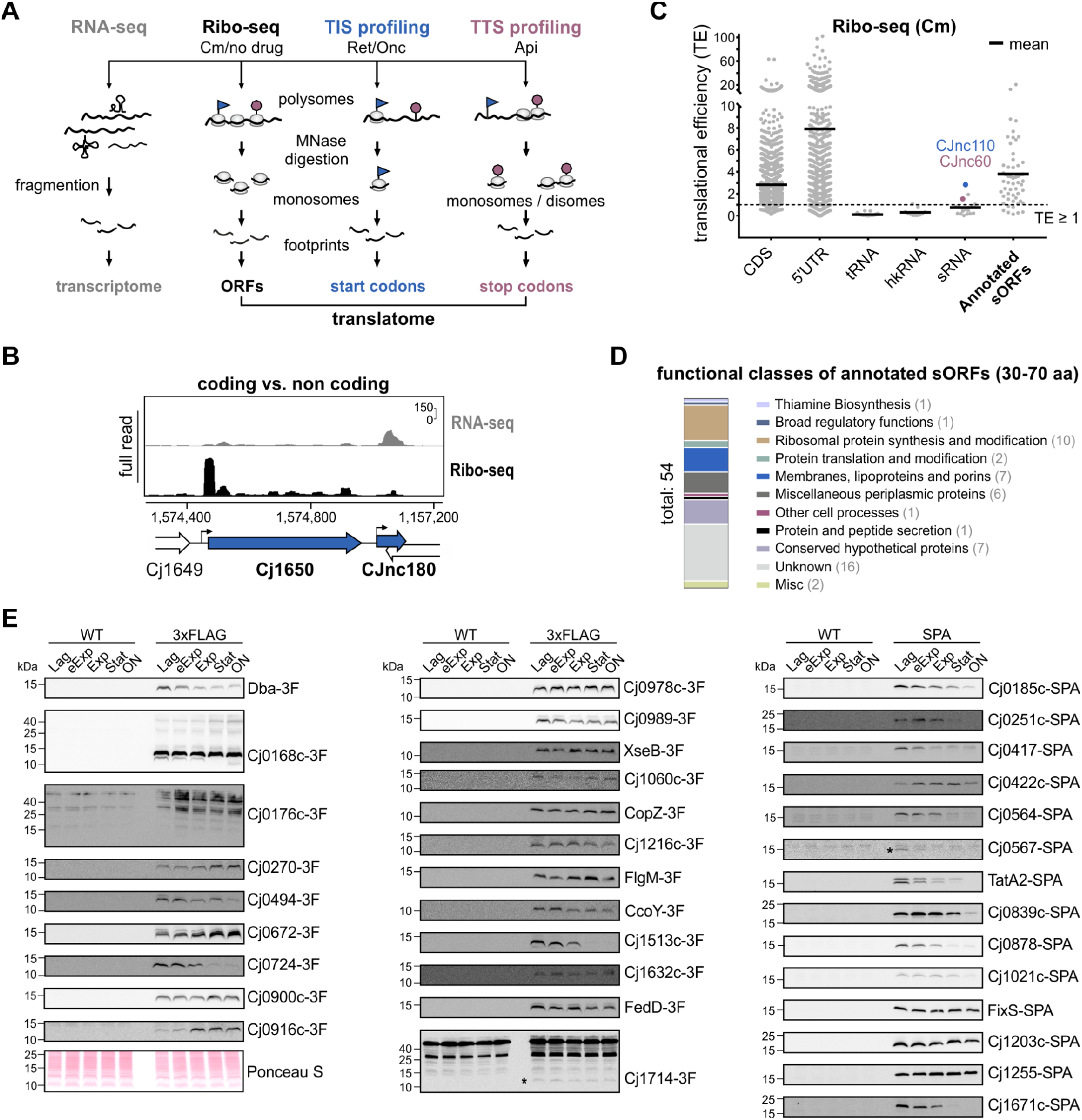
Establishing Ribo-seq in *C. jejuni* with the annotated small proteome. **(A)** Overview of Ribo-seq techniques applied/developed in this study. MNase: micrococcal nuclease, Cm: chloramphenicol, Onc: oncocin, Ret: retapamulin, Api: apidaecin. **(B)** Ribo-seq and paired RNA-seq distinguish the coding gene Cj1650 from the non-coding sRNA CJnc180 (Svensson and Sharma, 2021). Y-axis: rpm (reads per million). Representative of three independent experiments. **(C)** Translational efficiency (TE: Ribo-seq/RNA-seq) from Cm-treated Ribo-seq for different feature classes in the *C. jejuni* annotation (Dugar et al., 2013; Gundogdu et al., 2007). CDS: coding sequences/ORFs. hkRNA: housekeeping RNAs. **(D)** Functional classes for annotated sORFs (Gundogdu et al., 2007). See also **Table S1. (E)** Translation validation by WB for annotated sORFs with C-terminal 3xFLAG (3F) or SPA epitope tags with an anti-FLAG antibody. Ponceau S staining of membranes was used as loading control. eExp: early exponential. Stat: stationary. ON: overnight. Untagged WT: antibody (anti-FLAG) control. Representative of at least two independent experiments. Asterisks: low abundance FLAG-specific bands.

We first established canonical Ribo-seq experimental and data analysis protocols (with Cm, (Becker et al., 2013)) and adapted them for use in *C. jejuni* (**Fig. S1A**, see **Methods**). In general, our “standard” *C. jejuni* Ribo-seq dataset successfully differentiated between known coding and non-coding regions. For example, cDNA reads for the sRNA CJnc180 (Dugar et al., 2013; Svensson and Sharma, 2022, 2021) were mainly restricted to the RNA-seq (transcriptome) library prepared in parallel, while the adjacent ORF Cj1650 had both RNA-seq and Ribo-seq coverage (**Fig. 1B**). Housekeeping ncRNAs (hkRNAs) and tRNAs likewise showed low footprint coverage (*e*.*g*., the RNA component of the signal recognition particle) (**Fig. S1B**).

Global examination of translational efficiencies (TE: ratio of coverage in Ribo-seq/RNA-seq libraries) also showed a clear difference between coding and non-coding features, with ORFs having a mean TE ≥1 and tRNAs/housekeeping RNAs <1 (**Fig. 1C**). The TE of mRNA leaders was generally ≥1 and on average higher than coding regions. This might reflect the short length of *C. jejuni* 5′UTRs (median ∼25 nt), which means a significant length is protected from digestion by initiating ribosomes (Dugar et al., 2013; Porcelli et al., 2013). Also, re-initiation can occur in the absence of elongation, especially upon Cm treatment (Mohammad et al., 2019). Nonetheless, long annotated 5′UTRs showed absence of reads in Ribo-seq but not RNA-seq libraries far upstream of the start codon (*e*.*g*., Cj0978c, ∼194 nt; **Fig. S1C**). Most annotated sRNAs had a TE <1, indicating they are in fact non-coding. However, two intergenic sRNAs had a TE ≥1 (CJnc60 & CJnc110; 1.54 & 2.83, respectively), suggesting they might be small mRNAs or dual-function sRNAs encoding an sORF (**Fig. 1C**). In line with this, it has been proposed that CJnc60 encodes an unannotated SelW homolog (81 aa) (Shaw et al., 2012). Based on this successful set-up of Ribo-seq in *C. jejuni*, we next inspected translation of annotated small proteins in more detail.

### Analysis of translation and conservation of the annotated *C. jejuni* small proteome

The current *C. jejuni* NCTC11168 annotation (from 2021-09-11) includes only 54 small proteins (here based on a definition of ≤70 aa) and none below 30 aa (**Table S1**). These 54 small proteins include ten small ribosomal proteins (r-proteins) and a handful with annotated housekeeping functions (*e*.*g*., Dba, SecE), but mostly hypothetical and unvalidated proteins of unknown function (**Fig. 1D, Table S1**). Most had a TE >1 (mean ∼4) in our dataset, supporting that they are indeed translated (**Fig. 1C**). However, Ribo-seq can be prone to false positives due to, *e*.*g*., ribosome association without translation (Weaver et al., 2019). To further validate translation of these small proteins, as well as to guide sORF predictions based on our Ribo-seq data (below), we next generated in-frame epitope-tagged versions of 41 of all 44 annotated non-ribosomal sORFs at their C-terminus with either 3×FLAG or SPA (Sequential Peptide Affinity). For three annotated sORFs, we could not generate a tagged version at the native locus, suggesting that the epitope interfered with an essential function (Cj1047c/63 aa, Cj1160c/59 aa, *secE*/59 aa). We also included Cj0185c (69 aa), which had been removed from the current NCBI annotation (2021-09-11) but caught our eye during initial inspection of sORF predictions, as the prediction was located between two genes with a gap in locus tag numbering (**Fig. S1D**).

WB blotting detected 35 out of 42 tagged small proteins in at least one condition, and 34 in the same growth phase as Ribo-seq (log) (**Fig. 1E, Table S1**). This analysis also showed a diversity of expression patterns for the tagged small proteins. Although most showed stable or decreased levels at later growth phases, some accumulated at least two-fold as cultures progressed to stationary phase, suggesting they might mediate adaptation under these conditions (Cj0672, Cj0916c, CopZ, **Fig. 1E**). Independent MS analysis of whole cell protein samples also validated translation of 15/55 sORFs, including six small ribosomal proteins, as well as Cj0185c and untaggable Cj1047c (**Table S2**). Two or more unique peptides were detected for the majority of these.

We also inspected conservation of the annotated sORFs in Epsilonproteobacteria. Annotation-independent homology searches (tblastn, **Fig. S2**) showed strong conservation of several small proteins suggesting they might have important functions, including some with only hypothetical annotations. This included Cj0270 (tautomerase), as well as the aforementioned lipoprotein Cj0978c (**Fig. S1C**). Others were strain-specific, such as the putative transcriptional regulator Cj0422c (absent in widely-studied strain 81-176), or species-specific, such as Cj0168c (absent outside of *C. jejuni*). Previously-missing Cj0185c is conserved as far as *C. upsaliensis*, and includes a PhnA-domain of unknown function that is widely conserved in bacteria (**Fig. S1D**), while Cj0916c is a predicted selenoprotein (Cravedi et al., 2015). Small proteins are often membrane-associated (Orr et al., 2020; Yadavalli and Yuan, 2022). PSORTb localization predictions indicated that seventeen annotated *C. jejuni* small proteins have putative transmembrane helices, six encode possible signal peptides, and nine have an overall prediction of localization at the cytoplasmic membrane (**Table S1**) (Yu et al., 2010).

Overall, our inspection of expression, conservation, and sequence features suggested that *C. jejuni* encodes many *bona fide* small proteins with unexplored, hypothetical functions, which are potentially secreted or associated with the cell envelope. Our Ribo-seq and complementary WB/MS analysis suggests that the majority of annotated *C. jejuni* sORFs are translated in log phase and confirms that Ribo-seq is a sensitive approach for detecting translated small proteins. As the annotated small proteome most likely underrepresents the complete small proteome of *C. jejuni*, especially at the length range <30 aa and overlapping annotated ORFs, this systematically validated annotated sORF set provided a benchmark to guide novel sORF detection by Ribo-seq, TIS, and TTS (below).

### Retapamulin-based translation initiation site (TIS) profiling reveals start codons

Assigning Ribo-seq coverage to sORFs residing internal to or overlapping with longer ORFs can be challenging. To globally map ORFs, including any (s)ORFs internal to other ORFs with high confidence, we next established TIS profiling in *C. jejuni* to reveal start codons (**Fig. 1A**). We first aimed to apply Ribo-RET using Ret to stall initiating ribosomes (Meydan et al., 2019). However, Multi-drug efflux pumps of Gram-negative bacteria, such as AcrB-TolC, can reduce sensitivity to Ret. A Δ*tolC* mutant strain was used previously in *E. coli* to reduce efflux (Meydan et al., 2019). We found that deletion of the gene encoding the major efflux component CmeB in *C. jejuni* (Pumbwe and Piddock, 2002) reduced the Ret minimum inhibitory concentration (MIC) 8-fold (see **Supplementary Methods**). Moreover, treatment of the Δ*cmeB* mutant, which grew similarly to the WT strain under standard conditions (**Fig. S3A**) with 12.5 μg/ml Ret for 10 minutes collapsed polysomes to monosomes even without micrococcal nuclease (MNase) digestion (**Fig. S3B**), as reported for *E. coli* (Meydan et al., 2019). This indicated successful stalling of initiating ribosomes and run-off of elongating ribosomes. We omitted Cm treatment of the control/no drug culture for the TIS experiment to avoid potential bias in overall Ribo-seq coverage (Mohammad et al., 2019) and used a fast-filtration method to harvest cells (see **Methods**). Even without Cm, we still successfully captured polysomes and observed general enrichment of coding compared to non-coding features (**Figs. S3B & S3C**). This suggests that Ribo-seq can be applied without Cm, and we omitted it from all future experiments.

To determine whether Ret treatment in *C. jejuni* successfully enriched ribosome occupancy at start codons on a genome-wide scale, we performed metagene analysis of cDNA coverage near all annotated start codons (ATG, TTG, GTG). This showed enrichment in the Ret vs. no drug libraries at position -16/+16 nt upstream/downstream of start codons for 5′/3′ read end coverage, respectively (**Fig. 2A; Figs. S3D & S3E**). As reported for MNase-generated Ribo-seq libraries in other bacteria, trimming at 3′ ends was sharper than at 5′ ends (Mohammad et al., 2019) (**Figs. S3D & S3E**). Imprecise trimming by MNase in bacterial Ribo-seq can blur P-site offsets (*i*.*e*., the distance from the ribosome P-site/start codon to the trimmed read 3’ end), but can be improved by first binning reads based on their length and selecting those giving the best resolution (Bartholomäus et al., 2021; Mohammad et al., 2019). We used a similar approach, which revealed that while our protocol is designed to recover RNAs between 26-34 nt (for monosome footprints) (Ingolia et al., 2012), enrichment at TIS in our dataset was strongest for 31/32 nt-long reads (3′ end coverage).

**Figure 2.**
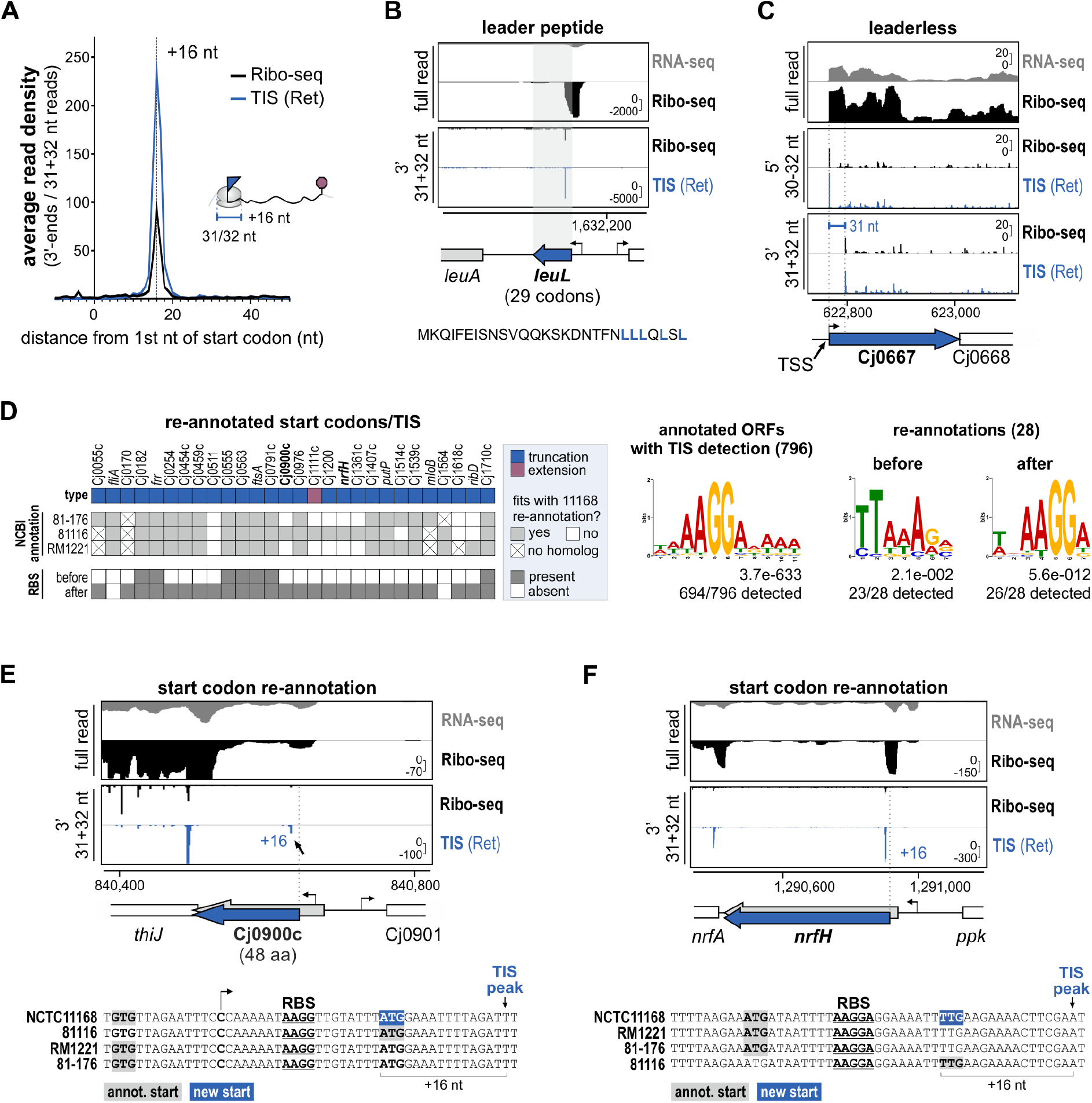
Ribo-seq & TIS-based mapping of start codons. **(A)** Genome-wide ribosome occupancy at start codons for no drug (Ribo-seq) and TIS(Ret) libraries. Representative of three independent experiments. See also **Figs. S3D & S3E. (B)** LeuL leader peptide (CJsORF41, **Table S1**). *Below*: proposed amino acid sequence (Porcelli et al., 2013). **(C)** Leaderless ORF Cj0667 (Porcelli et al., 2013). Leaderless ORFs are expected to have 5′/3′ peaks at the TSS/30 nt downstream of the TSS, respectively. **(D)** Start codon re-annotations based on Ribo-seq and TIS (see also **Table S4**). *Right side:* RBS motif predictions. *Left*: *C. jejuni* consensus RBS motif identified in 15 nt upstream of all annotated ORFs with a detected TIS.. *Center & right*: RBS motif detected upstream of 28 re-annotated start codons only after modification. Motifs were predicted with MEME (Bailey et al., 2009). **(E)** Re-annotation of Cj0900c sORF start codon. Grey: Annotation (59 aa) (Gundogdu et al., 2007). Blue: Re-annotation (48 aa). **(F)** Re-annotation of the *nrfH* start codon. *Bottom*: alignment of translation initiation region from four *C. jejuni* strains. Grey - annotated start codons. Blue - new NCTC11168 start codon. Underlined - potential RBS. Bent arrows - TSS (Dugar et al., 2013). Arrow - Ret-enriched peak. For screenshots, y-axis: rpm (reads per million).

Manual inspection of annotated ORFs using these optimal read-lengths and offsets revealed TIS signals at the expected position. For instance, at a highly translated, conserved ribosomal protein (r-protein) operon, we detected TIS vs. Ribo-seq enrichment at ∼16 nt downstream of each start codon (**Fig. S4A**). We also observed strongly enriched TIS peaks for three previously predicted but unannotated *C. jejuni* leader peptides of amino acid biosynthesis operons (Porcelli et al., 2013), including LeuL, as well as TrpL and MetL (**Figs. 2B & S4B**). Ret treatment also revealed a putative internal TIS in *lysC* driving translation of a LysCβ subunit, in agreement with the short isoform previously reported in *B. subtilis* (**Fig. S4C**) (Chen et al., 1987). We also validated translation of seven leaderless genes, such as Cj0667 (**Fig. 2C; Table S3**). For Cj0459c, which was previously suggested to be transcribed as a leaderless mRNA (Porcelli et al., 2013), our TIS data rather indicate that it is encoded on a leadered mRNA and requires start codon re-annotation (**Fig. S4D**). The status of 14 leaderless ORFs remains unclear, due to insufficient Ribo-seq coverage and/or absence of TIS peaks (**Table S3**). This might be partially caused by technical limitations, as our size-selection approach discards short footprints (<26 nt), such as those expected from ribosomes at leaderless start codons (Gelsinger et al., 2020). Therefore, the true number of leaderless ORFs might be larger.

Overall, metagene analysis and single-gene inspection supported the successful establishment of TIS profiling and its potential to map start codons in *C. jejuni*.

### Translation initiation site signals from Ret treatment refine start codons

Accurate start codon information is crucial for identification of regulatory elements such as 5’UTRs or binding sites for sRNAs, but manually annotating them can be laborious. To allow for automated detection and annotation of start codons based on TIS data, we adapted an approach for genome-wide detection of TIS peaks for re-annotation of start codons and discovery of TIS associated with potential new sORFs (see **Methods** for details) (Gelsinger et al., 2020). In brief, we used the above-mentioned parameters (read-lengths and offsets) from metagene analysis (**Fig. S3E**) for peak detection. With this approach, we detected peaks for approx. 50% of annotated start codons in our TIS(Ret) dataset (**Fig. S4E**).

To re-annotate start codons based on our TIS data, we next inspected A) ORFs without a detected TIS(Ret) peak for the annotated start, but with an in-frame start codon in close proximity (<51 nt), as well as B) 126 ORFs previously marked for re-annotation based on their TSS position in four *C. jejuni* strains (Dugar et al., 2013). Based on Ribo-seq/TIS coverage, we re-annotated the start codon of 28 ORFs out of 1576 annotated protein-coding genes in strain NCTC11168 (**Fig. 2D**; **Tables S4 & S5**). Most of the changes resulted in shorter N-termini, and most were supported by previous length comparisons in four *C. jejuni* strains (Dugar et al., 2013). Investigation of the 15 nucleotides upstream of start codons before or after re-annotation with MEME (Bailey et al., 2009) predicted an RBS motif for 26 of the 28 re-annotated ORFs after re-annotation, while none had a predicted RBS before re-annotation (**Fig. 2D**, *right*).

The 28 re-annotated genes included sORF Cj0900c (annotated as 59 aa), where we mapped the start codon downstream to generate an even shorter protein (48 aa) (**Fig. 2E**). Our analysis also suggested several relatively well-characterized proteins have shorter N-termini, including FliA (motility anti-sigma factor, **Table S5**), as well as NrfH (nitrite reductase small subunit), where instead of a canonical ATG, a GTG appears to be used, truncating the N-terminus by 8 aa (**Fig. 2F**). The TIS analysis also revealed an ORF (in addition to Cj0185c, **Fig. S1D**) that should be added back to the NCBI annotation. Inspection of Cj0055c, marked for re-annotation, suggested not only that its start codon should be moved 36 nt downstream to generate a shorter ORF (263 vs. 275 codons), but also that adjacent coverage might arise from the previously-annotated ORF Cj0056c (**Table S5**). MS data revealed two N-terminal extensions (Cj0636 and Cj1253 (*pnp*)) (**Table S2 & S5**).

Altogether, these observations suggest the successful establishment of TIS profiling with Ret in a new organism, *C. jejuni*. Moreover, our TIS (and paired Ribo-seq) data allowed us to detect previously-predicted short leader peptides/upstream ORFs (uORFs), inspect several predicted leaderless mRNAs, and refine start codon coordinates for 28 ORFs based on experimental data. This shows the overall utility of TIS profiling for annotation of *C. jejuni* coding regions, including sORFs.

### Apidaecin treatment enriches for terminating monosome and disome footprints

While internal TIS are well known to generate shorter, functional proteins (Meydan et al., 2018), 3′ end truncations are less well-characterized. C-terminal truncations might arise via, *e*.*g*., point mutations, alternative decoding, or ribosomal frameshifting. These features are, in principle, not encoded in, or are not easily detected from, the reference genome and might be best identified experimentally. In addition, we hypothesized that detection of stop codons (TTS) might provide additional evidence for canonical (s)ORF translation. Thus, we established a new Ribo-seq-based TTS profiling method to globally map stop codons. Our approach uses the PrAMP Api to trap terminating ribosomes (Florin et al., 2017; Mangano et al., 2020) and enrich Ribo-seq coverage at stop codons (**Fig. 1A**).

Although some *C. jejuni* strains are not sensitive to natural apidaecin from honeybees (Casteels et al., 1994), our NCTC11168 WT isolate had a minimum inhibitory concentration (MIC) for synthetic apidaecin 137 (hereafter, Api) of ∼6.25 μM, which was dependent on Cj0182, encoding a protein with homology to the SbmA peptide transporter required for uptake of PrAMPs in *E. coli* (Weaver et al., 2019) (see **Supplementary Methods**). We treated a *C. jejuni* WT culture with ∼10× MIC Api (50 μM) for 10 min. In parallel, we treated a culture with the same concentration of the PrAMP Onc (MIC ∼6.25 μM, see Methods for details) for parallel TIS profiling. Like Ret, Onc collapsed polysomes into monosomes without MNase treatment (**Figs. 3A & S3B**). In contrast, polysomes from *C. jejuni* treated with Api showed MNase resistance (**Fig. 3A**, *right*). We hypothesize that this results from increased proximity of adjacent ribosomes that results from stalled terminating ribosomes. This might interfere with MNase cleavage as has been reported in eukaryotic ribosomes near stop codons and collision sites and is exploited for so-called “disome” profiling (Han et al., 2020; Meydan and Guydosh, 2020; Wu et al., 2020). We thus also prepared libraries from disome footprints (50-80 nt) for Api-treated cultures. In total, our TTS profiling experiment included five libraries: RNA-seq, untreated Ribo-seq, TIS profiling using oncocin [TIS(Onc)], and two TTS profiling libraries: monosome [TTS(Mono)] and disome [TTS(Di)] libraries (**Fig. 3A**, *bottom*).

**Figure 3.**
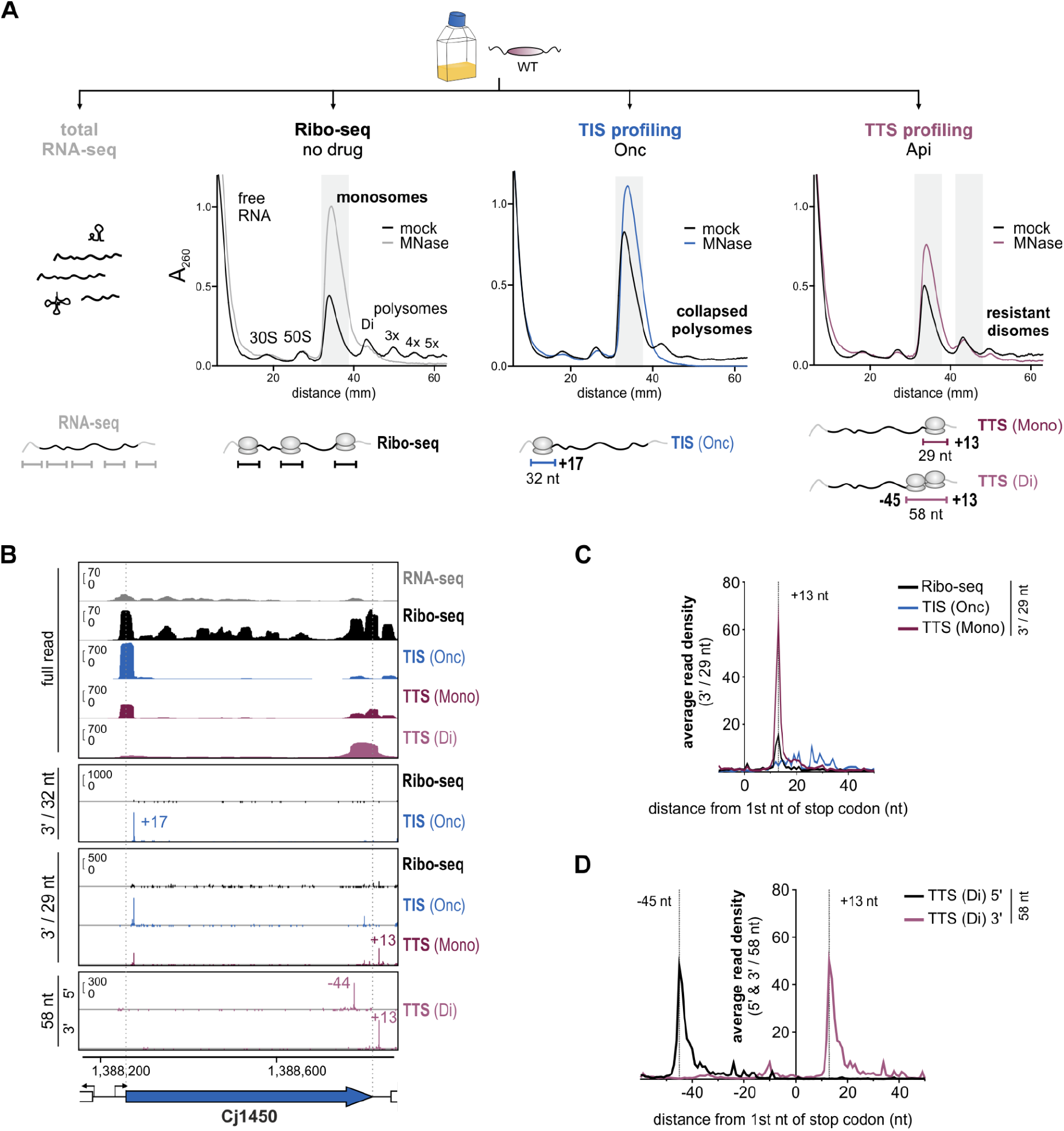
Apidaecin treatment combined with Ribo-seq for mapping translation termination sites. **(A)** Sucrose density gradient analysis of ribosome species in lysates for the TTS experiment for untreated (no drug), Onc (TIS), or Api (TTS) treated *C. jejuni* WT cultures. Shaded regions: fractions harvested for library preparation. Representative of three independent experiments. *Below:* Overview of libraries, footprint lengths, and 5′/3′ peak offsets used for TIS/TTS automated detection are shown (see also panels C/D). **(B)** Ribo-seq cDNA coverages (full-read and single-nt) for Cj1450. Single-nt coverage files (first or last base of reads) were generated with read lengths giving the strongest/sharpest enrichment/peaks (see panels C/D). Y-axis: rpm (reads per million). **(C & D)** Metagene analysis of ribosome occupancy at annotated stop codons for 3’ ends of 29 nt reads for the monosome library or 5**′**/3**′** ends of 58 nt reads for the disome library. All read lengths: **Fig. S5**. -45 nt/+13 nt: offsets used for TTS(Di) and (Mono) peaks, respectively.

Full-read cDNA coverage plots for single ORFs revealed enrichment of coverage towards the 3′ end for both the TTS(Mono) and TTS(Di) libraries compared to no-drug or TIS(Onc), suggesting that Api was stalling terminating ribosomes (*e*.*g*., Cj1450) (**Fig. 3B**). Increased coverage extending several footprint lengths before the Cj1450 stop codon suggested that Api was also inducing ribosome queuing. To determine genome-wide if Api-treatment enriched ribosome occupancy near TTS, we performed metagene analysis of coverage near annotated stop codons. This showed enrichment for 5′/3′ read end positions at -16 nt/+13 nt, respectively, in the TTS(Mono) vs. no drug libraries, which was most pronounced for 29 nt reads (**Figs. 3C & S5A**). Onc enriched 3′ read-end coverage vs. the no drug library at ∼17 nt downstream of start codons, and 32 nt reads produced the sharpest enrichment (**Figs. S5B & S5C**). Enrichment at +17 nt vs. start codons for Onc contrasted with enrichment at +13 nt vs. stop codons for Api (**Figs. S3E, S5A, S5C**). This 3 nt difference in peak offset at start vs. stop codons is in line with observations in *E. coli*, which show that Api leaves the stop codon in the A-site, while initiation inhibitors stall the ribosome with the start codon in the P-site (Florin et al., 2017; Meydan et al., 2019). For TTS(Di) libraries, peaks of ribosome occupancy near stop codons were highest at - 45/+13 nt for 5′/3′ end mapping, respectively (**Fig. 3D**). Peaks were highest and sharpest with 58 nt reads for both mapping approaches, in line with a disome footprint length (**Fig. S5D**).

We manually inspected coverage for 3’-end mapped data, restricted to read lengths that were suggested by above metagene analysis to give the sharpest enrichment (**Figs. S5A & S5D**), for a highly translated r-protein operon. This showed strongly enriched peaks in the TTS(Mono) vs. conventional Ribo-seq library at the expected +13 nt position downstream of each stop codon (**Fig. S6A**). Inspection of TTS(Di) coverage likewise revealed peaks at positions in line with metagene analysis (approximately - 45/+13 nt for 5’/3’-end mapped data, respectively; **Figs. S5D & S6A**). We did observe some artifacts of Api treatment previously observed in *E. coli* (Mangano et al., 2020), including increased coverage in some 3′UTRs, presumably caused by readthrough of stop codons upon Api treatment (**Fig. S6B**). This was observed for both TSS(Mono) and TTS(Di) libraries. We also observed enrichment at some start codons in TTS(Mono) libraries (**Figs. 3B & S5B**). Our unique parallel-generated Onc library therefore served not only to map start codons in WT, but also to control for this artifact. Start codon enrichment was also less pronounced for the TTS(Di) compared to the TTS(Mono) library, suggesting disome profiling might also circumvent some artifacts induced by Api. Overall, this suggests that despite some complexity around stop codons, Api treatment paired with Ribo-seq reveals sites of active translation termination.

### TTS profiling reveals stop codon usage not apparent from the reference genome

To demonstrate the utility of Api treatment in discovering ORFs based on their stop codons, we inspected several previously described/predicted *C. jejuni* genes that have TTS that would not be immediately apparent from the reference genome. This included those with stop codon recoding, pseudogenes generated by single point mutations, and phase-variable genes that rapidly change their translation status based on alterations at simple-sequence repeats. For example, the predicted *selW* ORF in *C. jejuni* in CJnc60 encodes a selenoprotein where a specific UGA codon is decoded by tRNA-SeC to incorporate a selenocysteine residue (Shaw et al., 2012). The full-length *selW* ORF is not immediately apparent as the selenocysteine-encoding UGA is interpreted as a stop codon. In line with full translation of an 80 aa SelW homolog, Ribo-seq revealed ribosome occupancy across the CJnc60 transcript, although this was uneven (**Fig. 4A**). Consistent with full translation, a TTS peak was present after the putative full-length *selW* stop codon and several MS peptides, detected by MS/MS, mapped beyond the putative tRNA-Sec-decoded UGA stop codon (**Table S2**).

**Figure 4.**
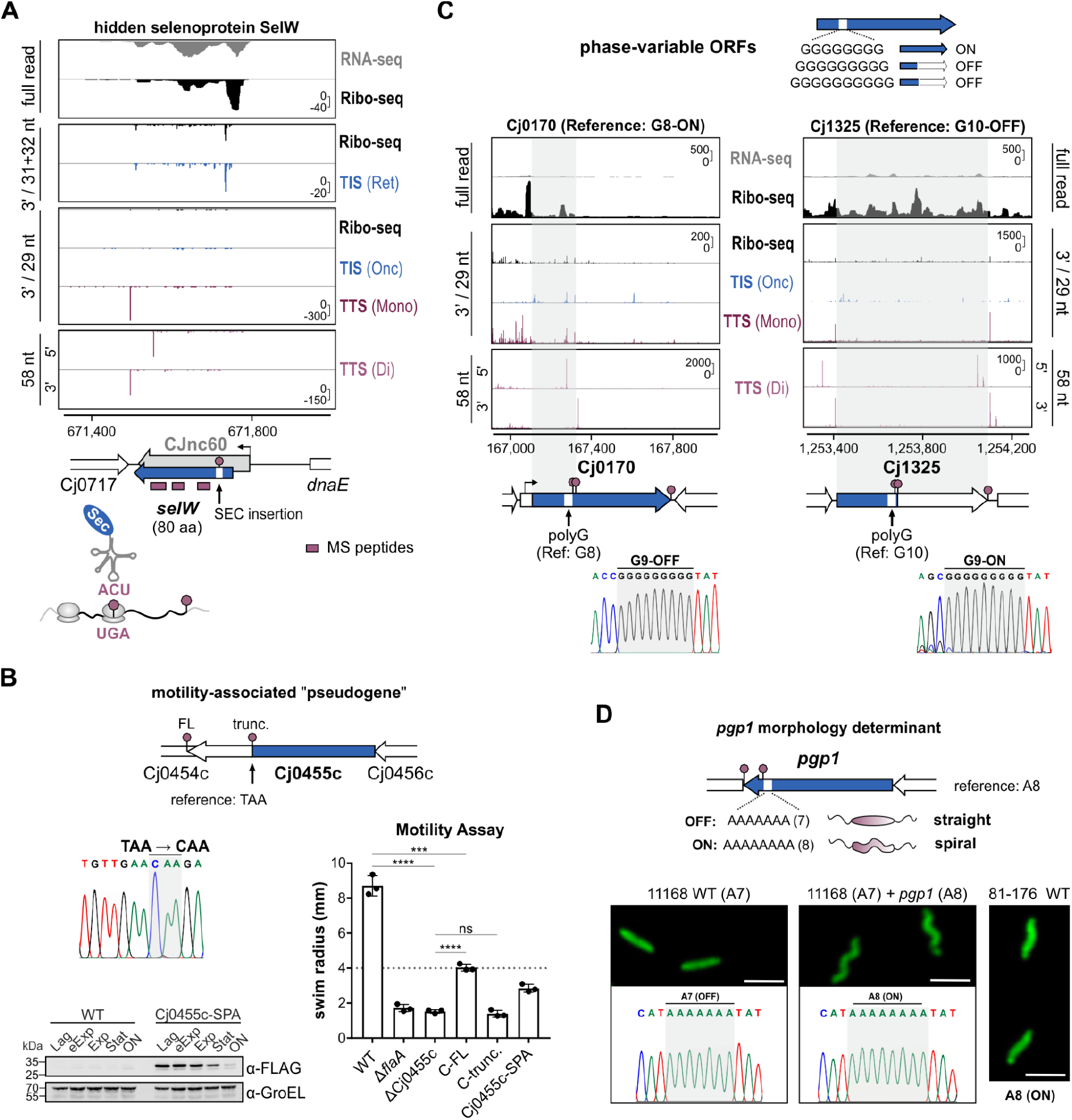
TTS profiling reveals translation status of infection-relevant genes not apparent from the reference genome. **(A)** Translation of the selenoprotein *selW*. Grey: CJnc60 sRNA (Dugar et al., 2013). Blue: *selW* ORF (Shaw et al., 2012). Red: detected MS peptides. Stop sign: UGA read by tRNA-Sec. **(B)** Full-length (FL) translation of motility-related “pseudogene” Cj0455c. *Top*: Genomic context with truncation due to premature TAA. *Bottom left*: TAA-CAA conversion (Sanger sequencing) and WB validation of FL translation (anti-FLAG). Representative of two independent experiments. *Bottom right*: Motility assay on soft agar of ΔCj0455c mutant complemented either with FL (C-FL; CAA) or truncated (C-trunc.; TAA) Cj0455c isoform at unrelated *rdxA*. Mean/standard deviation of swim radii for three biological replicates is shown. Δ*flaA*: non-motile control. ****: p <0.0001, ***: p <0.001, ns: p >0.05 (not significant), Student’s t-test. See also **Figs. S7A & S7B. (C)** TTS peaks for phase variable genes Cj0170 and Cj1325. Blue arrows: current NCBI gene annotation. *Below*: Sanger chromatograms of G-tract length in WT used for Ribo-seq/TTS. **(D)** *pgp1* shape determinant TTS peaks, genotype and morphology of WT isolate used for Ribo-seq. *Top*: genomic context of variable *pgp1* A-tract. A8: ON, A7: OFF/truncated. *Bottom*: Introduction of *pgp1*-A8(ON) allele into unrelated *rdxA* of rod-shaped WT restores spiral morphology. Cells were stained with FITC (fluorescein isothiocyanate) and imaged by confocal microscopy. See also **Figs. S7C-E**. Y-axis: rpm (reads per million).

We next inspected a list of *C. jejuni* pseudogenes (Gundogdu et al., 2007), which can be generated by single nucleotide changes that might be variable between strains, or generated by sequencing errors. Consistent with their genotype, several showed coverage only until the annotated premature stop codon as well as an associated TTS peak (**Fig. S6C & S6D**). In fact, some generated potential novel sORFs (*e*.*g*., derived from Cj0565) (**Fig. S6D**). In contrast, coverage for others appeared different from what would be expected from their annotation. For instance, Ribo-seq coverage for pseudogene Cj0455c extended beyond an annotated, premature in-frame TAA stop codon, while the TTS(Mono) library showed a peak downstream of a potential full-length ORF stop codon, rather than at the annotated premature stop codon (**Figs. 4B & S7A**). In line with this observation, a 61 aa C-terminal extension was previously reported in some *C. jejuni* NCTC11168 isolates (Frirdich et al., 2017), and Sanger sequencing of our strain showed a similar TAA→CAA conversion.

To demonstrate that the full-length Cj0455c protein is in fact translated, we introduced a SPA epitope in-frame at the penultimate codon of the full-length ORF at the native locus. WB analysis detected translation of a protein of the expected full-length size (20.6 kDa + SPA tag) (**Fig. 4B**, *bottom*). Cj0455c was previously suggested to support *C. jejuni* motility, a key virulence factor in this pathogen (Gao et al., 2017). To provide additional evidence for the full-length genotype and the utility of TTS profiling in revealing translation not apparent from the reference sequence, we deleted Cj0455c and measured motility. This resulted in a non-motile phenotype compared to the parental strain. Introduction of a full-length (CAA), but not truncated (TAA), copy of Cj0455c into this ΔCj0455c strain at the unrelated *rdxA* locus partially restored motility, confirming that the phenotype of the deletion is not due to polar effects (**Figs. 4B**, *right* **& S7B**). Altogether, this suggests Cj0455c should be re-annotated as an intact ORF and confirms that Cj0455c encodes a motility-associated protein.

### TTS-profiling detects the ON/OFF status of phase-variable genes

*C. jejuni* regulates several ORFs via ON/OFF phase variation due to frameshifting at length variation of hypermutable homopolymeric tracts (*e*.*g*., polyG, polyA) (Burnham and Hendrixson, 2018; Parkhill et al., 2000). While most known phase-variable ORFs had low read coverage in our data, several had sufficient expression to reveal TTS peaks in line with their reference genome homopolymeric tract length. However, Ribo-seq coverage for two examples, previously found to be variable in human challenge experiments (Crofts et al., 2018), did not match their genotype. Reads for Cj0170 (reference G8, corresponding to the ON status) covered only the first half of the ORF until just after the G-tract (**Fig. 4C**, *left*). In line with the presence of a premature in-frame stop codon due to a G9 tract, a TTS(Di) peak was also visible just downstream of the polyG sequence. Sanger sequencing of genomic DNA confirmed that the dominant genotype of our WT isolate was G9-OFF instead of G8-ON. Paired TIS(Onc) data also showed that the Cj0170 start codon should be re-annotated downstream of its TSS, as suggested previously (Dugar et al., 2013) (**Table S4**). Translatome coverage for a second phase-variable ORF, Cj1325, was also inconsistent with its reference G-tract length (G10-OFF). Ribo-seq coverage extended across the entire Cj1325 ORF, and peaks were detected in the Api-treated libraries immediately downstream of the full-length stop codon (**Fig. 4C**, *right*). Consistent with Ribo-seq and TTS, Sanger sequencing showed that our isolate carried a G9-ON sequence.

*C. jejuni*’s characteristic helical cell shape is also under phase variable control, with one of the most common changes associated with rod morphology being a single nt conversion in a mutable A-stretch (A8-A7) in *pgp1* (Cj1345c, peptidoglycan endopeptidase) (Esson et al., 2016). Variation in this A-tract can generate a premature stop codon that truncates 14% of the C-terminus of the protein to render it non-functional (**Fig. S7C**). Our NCTC11168 isolate displays straight morphology (Dugar et al., 2016). TTS(Mono) and (Di) coverage showed peaks before the 3’ end of the *pgp1* coding region (**Figs. 4D** (*top*) **& S7D**). Closer inspection showed that the peaks were 13 nt downstream of a premature in-frame stop codon for the A7-OFF allele, and Sanger sequencing confirmed that our isolate harbors an A7 tract. Finally, to demonstrate again that TTS profiling can reveal physiologically relevant genomic changes that alter coding potential, we added a second A8-ON allele of *pgp1* (from the spiral strain 81-176) to our NCTC11168 WT isolate at the unrelated *rdxA* locus. This restored our isolate to spiral morphology (**Figs. 4D** (*bottom*) **& S7E**). Altogether, inspection of *C. jejuni* ORFs that can undergo relatively frequent changes in translation due to single nucleotide variations from the reference sequence in our novel TTS dataset demonstrated the potential of TTS profiling to inform on annotation-independent translation status.

### Ribo-seq mediated discovery of novel sORFs in diverse genomic contexts

Finally, we generated novel sORF predictions from all of our datasets. For predictions using classical Ribo-seq, we used two published tools in our HRIBO pipeline: REPARATION and DeepRibo (Clauwaert et al., 2019; Gelhausen et al., 2021; Ndah et al., 2017). As these tools can generate long lists of candidates with little guidance for ranking and many are likely false positives (Gelhausen et al., 2022), we used our WB/MS-validated small proteins (**Figs. 1E & S2**) as a ground-truth set to benchmark cutoffs for RNA-seq RPKM, TE, as well as the score generated by DeepRibo (**Fig. S8A**, see Methods for details). With relatively stringent cutoffs, conventional Ribo-seq based predictions recovered 39/45 validated sORFs (**Fig. S2**). In contrast, most ncRNAs were not detected (*e*.*g*., sRNA CJnc180 and signal recognition particle RNA (**Figs. 1B & S1B**). We also used the validated sORF set, most of which had a detected TIS (**Fig. S2**), to filter predictions from our TIS peak detection tool for strong candidates based on peak height and TIS/Ribo-seq fold-change cutoffs (**Fig. S8B**, see **Methods**). The peak detection tool was also used to predict sORFs associated with TTS (disome and monosome) peaks, with minor modifications (see **Methods**). Of the validated sORF set, 33/45 were detected by this approach in the monosome and/or disome libraries, and TTS(Mono)- and TTS(Di)-based predictions were mostly complementary (**Figs. S2 & S8C**). Overall, our systematically validated annotated sORF set indicated that the tools can identify translated sORFs from our data with both sensitivity and specificity (**Fig. 1E & Fig. S2**) and provided a resource to guide subsequent detection of novel sORFs.

With the tools and parameters described above, we compiled a filtered list of 421 *C. jejuni* sORF candidates of 11-71 codons in length from the predictions generated from our three Ribo-seq datasets (**Fig. 5A**) (see Methods & **Fig. S8D** for filtering details). Additional manual inspection of the top 16 TIS candidates in the range of 4-10 codons was included. Candidates from automated predictions included several likely positive unannotated sORFs that we observed during manual inspection, supporting their validity. For instance, the list included several short potential sORFs predicted based on Ribo-seq, TIS, and/or TTS, on CJnc60 that overlapped *C. jejuni selW* (80 aa) (**Fig. 4A**). As mentioned above, translation of *selW* depends on read-through of a stop codon by a special tRNA (Shaw et al., 2012), explaining why full-length *selW* was not predicted. The three uORFs (*leuL, trpL*, and *metL*) (**Figs. 2B & S4B**) were also included in filtered HRIBO, TIS, and/or TTS predictions, while TIS predictions captured the re-annotated version of the Cj0900c sORF (**Fig. 2E**).

**Figure 5.**
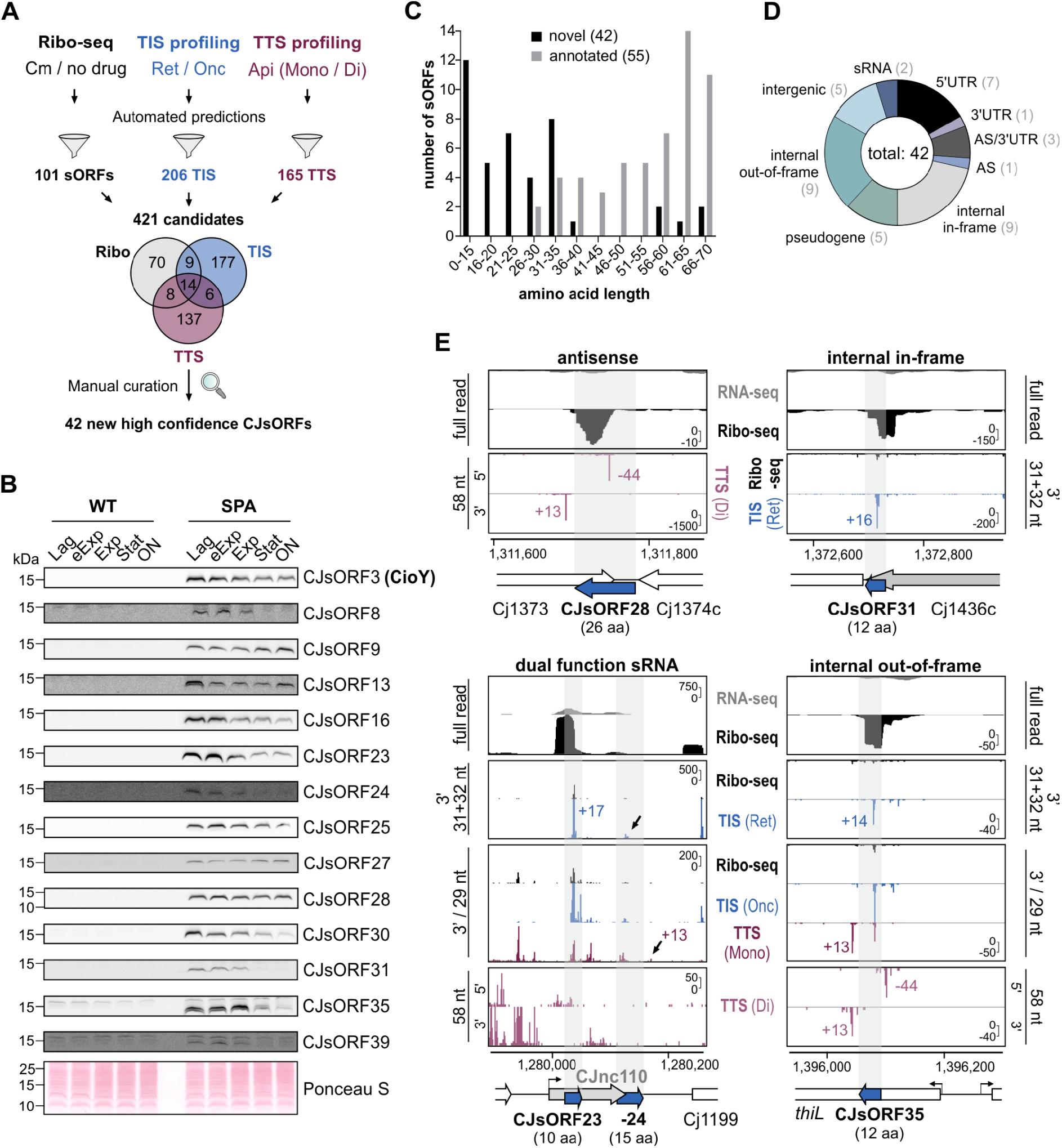
Novel *C. jejuni* sORFs revealed by integrated translatomics. **(A)** Workflow for expanding the *C. jejuni* small proteome via automated predictions based on ribosome occupancy, TIS, and TTS. Predictions were filtered (see Methods & **Fig. S8D**) for a set of 421 strong candidates, which were manually inspected for Ribo-seq, TIS, and TTS coverage & RBS. **(B)** WB validation of selected novel CJsORFs with a C-terminal SPA tag on Tricine-SDS-PAGE gels. SPA-tagged proteins were detected with an anti-FLAG antibody. Ponceau S staining of membranes served as a loading control and is shown only once. Representative of at least two independent experiments. eExp: early exponential. Exp: exponential. Stat: stationary. ON: overnight. **(C)** Length distribution of novel vs. annotated CJsORFs. **(D)** Genomic context of novel CJsORFs. AS: antisense. **(E)** Coverage for selected validated novel CJsORFs in diverse contexts by Ribo-seq, TIS, and/or TTS. See also **Fig. S10**. Y-axis: rpm (reads per million).

We next curated the 421 candidates, generated with these automated predictions and cutoffs, manually in a genome browser based on their Ribo-seq, TIS, and TTS coverage. We considered the shape of Ribo-seq coverage with respect to the sORF prediction, TIS/TTS peak position and height with respect to the prediction start/stop codon, and enrichment (*e*.*g*., RNA-seq vs. Ribo-seq, TIS vs. Ribo-seq), as well as presence of a potential RBS or transcriptional start site. This left us with a list of 42 new high-confidence “CJsORFs”, numbered based on genome position in strain NCTC11168 (**Table S1**). None were detected in our parallel MS-based proteomics survey using small protein-targeted approaches (**Table S2**). However, 37/42 had an RBS-like sequence 15 nt upstream of their start codons, supporting that they are true sORFs (**Fig. S9A**). We selected 17 from diverse genomic contexts for validation by SPA tagging. WB analysis of lysates from tagged strains growing in rich medium showed that 14 are robustly translated in at least one growth phase (**Figs. 5B & S9B**). While many were constitutively expressed, others accumulated or decreased at least ∼2-fold between exponential and stationary phase (*e*.*g*., CJsORF9 (33 aa) or CJsORF3 (34 aa), respectively). Low levels of three potential internal in-frame candidates (CJsORF7, CJsORF18, CJsORF36) in addition to CJsORF31 were also detected (**Figs. S9D-F**). We cannot, however, rule out that detected bands represent C-terminal fragments of the parental protein, generated by proteolysis, which retain the epitope tag.

The 42 novel CJsORFs were in general shorter than those in the current annotation, including 28 new genes with a length of ≤30 codons and the shortest encoding a protein of only 4 aa in length (**Fig. 5C**). This included a putative highly expressed ORF upstream of Cj0772c (CJsORF19, 6 aa; **Fig. S10A**). The new CJsORFs were encoded in diverse genomic contexts, including in 5′/3′UTRs, on sRNAs, internal in ORFs, and antisense to ORFs (**Fig. 5D**). For example, we validated a 26 aa sORF (CJsORF28) in the 3′UTR of Cj1374c (purine NTP pyrophosphatase) overlapping the Cj1373 membrane protein gene on the antisense strand (**Figs. 5E** (*top left*) **& S10B**). Our data also revealed two sORFs (CJsORF23 and CJsORF24; 10 and 15 aa, respectively) on CJnc110 sRNA, which we could validate by WB (**Figs. 5B & 5E**, *bottom left*). Because we used both Ribo-seq and start/stop codon detection, we were able to detect sORFs hidden in the Ribo-seq coverage of longer genes. For example, TIS profiling revealed the internal in-frame CJsORF31, translated from an internal start codon in Cj1436c and comprising only the last 12 aa of the 390 aa parental ORF (**Figs. 5E** (*top right*) **& S10C**). Finally, all three Ribo-seq methods revealed even internal out of frame examples, such as CJsORF35 (12 aa) within the *thiL* gene, encoding thiamine monophosphate kinase (**Fig. 5E**, *bottom right*).

Altogether, our translatomics approach, supported by independent western blot validation, expanded the number of *C. jejuni* small proteins almost by a factor of two (from 54 to 97) and suggested that the components of the *C. jejuni* small proteome are even shorter than what was previously annotated.

### A new small protein component of the CioAB terminal oxidase

To examine the new CJsORF primary sequences for potential functions, we performed tBLASTn searches in selected Epsilonproteobacteria strains, with parameters used previously for bacterial sORFs (Allen et al., 2014). This showed that the most highly conserved CJsORFs were generally those internal in-frame in annotated proteins with core functions, such as *flgH* (validated CJsORF18, 20 aa) (**Figs. 6A & S11A**). However, several novel candidates not overlapping annotated ORFs in-frame showed also relatively high conservation (*e*.*g*., CJsORF3 (34 aa), CJsORF25 (69 aa), CJsORF30 (21 aa)). Others appeared to be encoded by only some strains, such as the validated CJsORF31 (12 aa), which is encoded internally in the strain-specific gene Cj1436c. We also used BLASTp analysis to determine if any novel CJsORF might be already annotated as a stand-alone gene outside of *C. jejuni* NCTC11168. Several are in fact included as hypothetical proteins in the annotations of other *C. jejuni* strains, such as CJsORF3, corresponding to C8J_0076 in *C. jejuni* strain 81116 or CJE0079 in strain RM1221, respectively (**Table S5**). CJsORF38 (57 aa) is even annotated as Cj1558 in other sequenced derivatives of *C. jejuni* NCTC11168 such as NCTC11168-BN148 and appears to be another example that is not included in the current NCBI annotation despite showing evidence for translation in our datasets.

**Figure 6.**
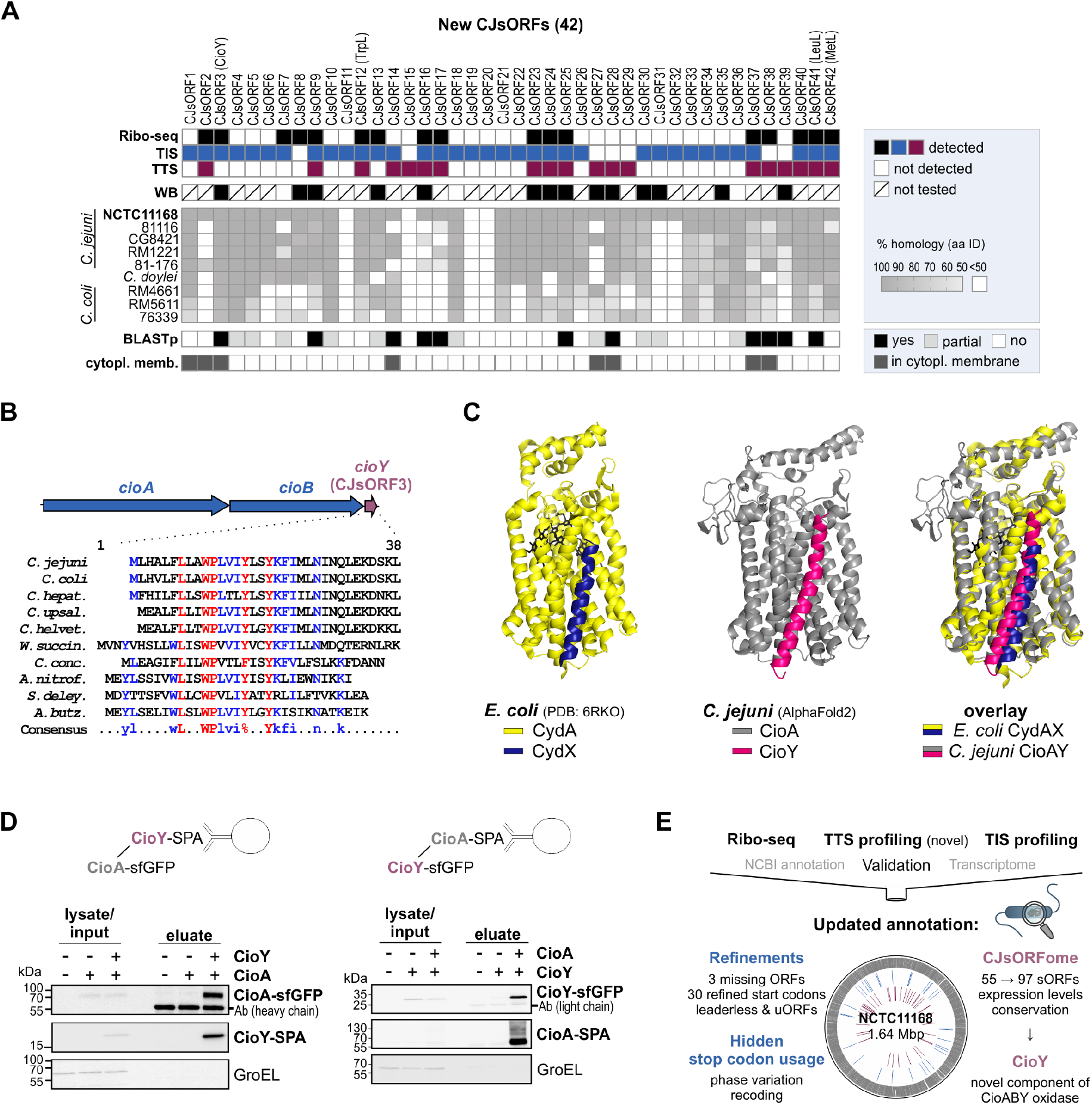
Integrated translatomics detects conserved *C. jejuni* small proteins with new functions. **(A)** Summary of detected novel CJsORFs along with conservation analysis. WB: detected by western blot at any growth phase tested. For conservation, sequences from *C. jejuni* NCTC11168 were used for tBLASTn or BLASTp searches at NCBI (see **Methods**). Cytopl. memb.: PSORTb predictions (Yu et al., 2010). See also **Fig. S11A. (B)** Genomic context and alignment of CioY sequences detected by tBLASTn in Epsilonproteobacteria. *C. jejuni*: NCTC11168. *C. coli*: RM4661. The alignment was generated using multalin (Corpet, 1988). **(C)** Predicted complex of *C. jejuni* CioA (grey) and CioY (magenta) using AlphaFold-multimer at ColabFold (Mirdita et al., 2022) compared to *E. coli* CydX (blue) with CydA (yellow; CydAX cryoelectron microscopy structure from (Safarian et al., 2019) (PDB 6RKO)). CydB and CydH subunits are not shown. **(D)** Reciprocal co-immunoprecipitation (coIP) from *C. jejuni* lysates reveals CioA-CioY interaction. CioY-/CioA-SPA were immunoprecipitated and detected by WB with an anti-FLAG antibody. sfGFP-tagged CioA/CioY were detected with anti-GFP. GroEL: loading control. Ab: heavy/light chain of anti-FLAG antibodies used for coIP. Representative of two independent experiments. **(E)** Overview of *C. jejuni* translatome refinements.

Subcellular localisation analysis using pSORTb (Yu et al., 2010) revealed that many of the novel CJsORFs have potential transmembrane helices (nine), signal peptide sequences (two), and/or are predicted to be localized to the cytoplasmic membrane (eight), suggesting they might be secreted or have functions at the cell envelope (Yu et al., 2010) (**Fig. 6A, Table S1**). This analysis again highlighted CJsORF3, annotated as a hypothetical protein outside of NCTC11168, which appears to consist mostly of a single transmembrane alpha helix. Alignment of potential CJsORF3 homologs from diverse Epsilonproteobacteria showed strong conservation of tryptophan, proline, and tyrosine residues (**Figs. 6B & S11A**). Moreover, we observed that all detected homologs were present downstream of components of a cytochrome *bd* terminal oxidase (in NCTC11168 annotated as *cydAB*) (**Fig. S11B**). Based on similarity to the *Pseudomonas* complex, the *C. jejuni* oxidase has been re-named CioAB (cyanide insensitive oxidase) (Jackson et al., 2007). *E. coli* CydA and CydB form a membrane-bound complex that includes two small proteins (CydX and CydH) (Safarian et al., 2019). Literature searches also revealed that CJsORF3 was previously identified by manual inspection (but not homology searches) of regions downstream of Epsilonproteobacterial *cydB* for potential *cydX* homologs and termed *cydY*, but was not validated or studied (Allen et al., 2014). Based on these observations taken together, we renamed the CJsORF3-encoded small protein CioY and further investigated its function in *C. jejuni*.

*E. coli* CydX binds CydA (**Fig. 6C**, *left*) (Safarian et al., 2019; VanOrsdel et al., 2013). To determine if *C. jejuni* CioY can interact with CioA, we performed structure and complex predictions with AlphaFold2-multimer at ColabFold (Mirdita et al., 2022). The results strongly supported that a CioY alpha helix interacts with CioA in a highly similar fashion as CydX-CydA (**Fig. 6C**, *centre & right*), despite limited sequence similarity of both small proteins (Allen et al., 2014). To validate that CioY interacts with CioA, we performed reciprocal co-immunoprecipitation (coIP) with differentially epitope-tagged small protein and potential interaction partner. WB analysis of eluates clearly showed that CioY binds CioA, suggesting that CioY is a new component of the CioAB terminal oxidase (**Fig. 6D**).

## DISCUSSION

Here, we have integrated several Ribo-seq approaches to map and annotate the protein coding potential genome-wide of the foodborne pathogen *C. jejuni*. In addition to doubling the *C. jejuni* small proteome, we also provide a comprehensive update of the *C. jejuni* ORF annotation (Gundogdu et al., 2007)(**Figs. 6E & S12A**). Our single-nt/codon resolution approach allowed us to validate leader peptides, leaderless ORFs and re-annotate start positions of genes with roles in key *C. jejuni* phenotypes (NrfH, FliA) (Hendrixson and DiRita, 2003; Pittman et al., 2007). Our study will facilitate identification of new physiological and regulatory mechanisms underlying *C. jejuni* survival and virulence.

### Termination site profiling detects stop codons in dynamic and hidden contexts

Besides establishing TIS-profiling in *C. jejuni*, we also developed a novel application of Api and disome profiling to map stop codons. *E. coli* Api Ribo-seq data, which were originally generated to study the mechanisms of translation inhibition by Api, were only recently reanalyzed for sORF detection (Mangano et al., 2020; Stringer et al., 2021). Besides sORF identification, we also demonstrated several additional uses of TTS profiling. The method revealed stop codons (or non-stop codons) that are absent, or are not immediately apparent, from the *C. jejuni* reference genome, including in phase variable ORFs, which are well-known to encode infection-related proteins in diverse pathogens (van der Woude and Bäumler, 2004). TTS profiling could reveal additional examples that are challenging to infer from the genome, such as those generated by ribosomal frameshifting. Recent MS-based proteomics analysis in *Salmonella* has led to the hypothesis that some pseudogenes, a hallmark of host-adapted strains, maintain full-length transcription and can likewise also show partial full-length translation, possibly via re-coding (via frameshifting or codon re-definition) mechanisms at or near the introduced nonsense mutation, to expand coding potential to provide opportunities in a more generalist niche (Feng et al., 2022). TTS profiling could, in principle, be applied to identify examples of such loci, or even - under relevant conditions - measure the ratio of full-length vs. prematurely terminated translation.

Our data suggest that TTS profiling can be a generic method that can be applied to diverse prokaryotes that are sensitive to Api to map stop codons. A requirement for application of Api to Gram-negative species, including *C. jejuni*, appears to be a homolog of the SmpB transporter, as reported for Onc (Weaver et al., 2019), although tightly-regulated heterologous induction of these PrAMPs might circumvent this (see below). Our study also suggests some additional guidelines for applying the approach meaningfully in other bacteria. As reported in *E. coli*, Api treatment introduced noise at *C. jejuni* TTS due to ribosome queuing (before) and readthrough (after). To circumvent this, we omitted stop codons <25 nt downstream of annotated TTS. Also as reported in *E. coli* and hypothesized to be due to lower affinity ribosome binding of Api even in the absence of RF1/RF2 (Mangano et al., 2020), Api also enriched ribosomes at some *C. jejuni* TIS. However, our parallel Onc library and disome approach (selective for ribosomes at TTS) mitigated some of the effects of this artifact. Disome footprints have been observed at stop codons in untreated eukaryotic cells (Han et al., 2020; Meydan and Guydosh, 2020), suggesting that disome profiling could be used in Api-insensitive species, although collisions at stop codons were rarely detected in untreated *E. coli* (Fujita et al., 2021).

### Complementary Ribo-seq approaches provide a more complete sORF census

Comparison of annotated sORF detection by the three methods (*i*.*e*., Ribo-seq, TIS, TTS) showed that a single approach alone was not sufficient to reveal all of our independently validated benchmark sORFs, which we used to guide our genome-wide automated predictions (**Fig. S2**). Based on this variable detection, we used more flexible criteria for our sORF predictions, and several of our 42 novel CJsORFs, including some validated by WB, were predicted from only a single dataset (five TIS-only, three TTS-only). A recent re-analysis of public *E. coli* TIS/TTS datasets required sORFs to have both start and stop codon signals (Stringer et al., 2021). Our results demonstrate that while using several Ribo-seq approaches and requiring signals in all datasets can increase confidence, more flexible criteria for sORF predictions might reveal additional, *bona fide* small proteins. Our approach also shows the utility of an experimentally validated sORF set to guide cutoffs to cope with high numbers of tool predictions (Gelhausen et al., 2022), and especially that manual inspection of Ribo-seq coverage and independent validation is an essential part of Ribo-seq.

To apply TIS and/or TTS profiling to other species, susceptibility to the used antibiotic or antimicrobial peptides is essential. For retapamulin sensitivity, an efflux pump mutant, such as the Δ*cmeB* strain used here or Δ*tolC* as used for *E. coli* (Meydan et al., 2019; Weaver et al., 2019) might be required for Gram-negative species. In contrast, for the uptake of PrAMPs such as Onc or Api, an SmbA peptide transporter or corresponding homolog might be required (Weaver et al., 2019). An alternative option for PrAMPs is *in vivo* expression via an available tightly regulated inducible system, as demonstrated to be feasible for Api in *E. coli* (Baliga et al., 2021).

Our Ribo-seq study, in addition to others, suggest that small proteomes in diverse bacteria and archaea are larger than what is currently annotated (Fuchs et al., 2021; Gelsinger et al., 2020; Laczkovich et al., 2022; Smith et al., 2022; Stringer et al., 2021; Venturini et al., 2020; Weaver et al., 2019). Taking a combined Ribo-seq approach, coupled with validation, is a strategy to generate a robust catalog of small proteins for future study.

### Function of *C. jejuni* sORFs and small proteins

Current understanding of bacterial sORF function places them into two categories: short proteins that interact with larger proteins/complexes to support/regulate their function, and short genes whose translation regulates adjacent ORFs. Ribo-seq revealed the small protein CioY, whose synteny suggested a function related to the CioAB terminal oxidase (Allen et al., 2014). CioY might support oxidase maturation or activity, for example by binding CioA and aiding assembly of/stabilizing the di-heme center as is the case for *E. coli* CydX/CydA (Hoeser et al., 2014; Safarian et al., 2019) despite limited homology. However, heme components of *C. jejuni* CioAB are unknown (Jackson et al., 2007). Terminal oxidases appear to be rich sources of small proteins (Garg et al., 2021; Koch et al., 2000; Kohlstaedt et al., 2017, 2016). Inspection of regions lacking annotated features adjacent to operons encoding large protein complexes is therefore likely a fruitful strategy for revealing new functional proteins in bacteria. Our set of 97 sORFs can now be inspected for roles in key adaptive strategies by measuring expression of our epitope-tagged versions under different conditions or by re-analysis of available datasets using our updated annotation as performed for *Salmonella* (Venturini et al., 2020).

While the current NCBI annotation for *C. jejuni* NCTC11168 includes no ORFs below 30 aa (Gundogdu et al., 2007), 28 of our 42 novel sORFs were ≤30 aa (shortest: 4 aa). The shortest functional bacterial small proteins so far include processed signaling peptides (5-10 aa), antimicrobial peptides such as microcin C (7 aa) (Severinov and Nair, 2012), and the *E. coli* CRP (cAMP receptor protein) regulating small protein SpfP (15 aa) encoded on sRNA Spot 42 (Aoyama et al., 2022; Neiditch et al., 2017). Another mode of action for many characterized sORFs is to regulate adjacent genes via their translation. In line with this, we identified the Met-codon enriched (MMYQMR) CJsORF19, a potentially new leader peptide that could regulate expression of the downstream Cj0772c (D-methionine transport system substrate-binding protein). Likewise, CJsORF15 (MAFY) is located upstream of a potential multidrug efflux protein (Cj0560), and could be related to translation-related control of expression of the downstream gene in the presence of translation inhibitors, as reported in, *e*.*g*., *Bacillus* (Takada et al., 2022).

At least 20 CJsORFs overlap annotated genes. Two CJsORFs overlap the infection-related sRNA CJnc110 (Kreuder et al., 2020). However, as regulatory targets for CJnc110 are so far not clear, CJnc110 might be a small mRNA, rather than a dual (regulatory and coding) function sRNA. While in-frame, ORF-internal start codons that generate protein isoforms are an accepted phenomenon in bacterial genome architecture, the function of internal out-of-frame ORFs is enigmatic (Meydan et al., 2018). CJsORF35 (12 aa), which is encoded out-of-frame in *thiL*, appears to be more highly expressed than the parental ORF and might be encoded on a separate ORF-internal transcript with an independent function, as recently shown for several *E. coli* genes *(Adams et al*., *2021)*. We also validated CJsORF28 (26 aa) opposite to integral membrane protein Cj1373. The function of antisense ORFs is mostly elusive (Zehentner et al., 2020). More generally, it is unclear how many sORFs are nonfunctional outcomes of pervasive translation, as many are not under purifying selection or do not show biochemical features, including stability, of *bona fide* proteins (Fijalkowski et al., 2022; Smith et al., 2022; Stringer et al., 2021). Nonetheless, pervasive translation might provide substrate for evolution of new small proteins (Hemm et al., 2020).

### Improving genome annotations to support study of bacterial pathogens

Recently, new *in silico* ORF prediction tools have been developed to identify small proteins in bacteria as well as phages based on genome sequence alone (Durrant and Bhatt, 2021; Fremin et al., 2022; Miravet-Verde et al., 2019). However, two examples (ranSEPS, smORFinder) did not detect most of our validated CJsORFs (**Table S1**). While laborious, experimental methods are still needed to annotate bacterial genomes, our validated sORF set could guide the development of the next generation of algorithms. The decreasing costs of deep sequencing means methods like Ribo-seq and TIS/TTS profiling, as well as genome-wide screens for gene function such as Tn-seq, can now be applied comparatively to diverse isolates and integrated with genomics data to complete annotations (Kobras et al., 2021).

Nonetheless, the rapidly expanding catalog of bacterial RNA-seq and Ribo-seq studies is raising important questions about how generated insights should best be used to inform annotations. How and when should formal annotations be adjusted? How should this information be curated? While these questions are considered, we have provided our datasets and annotation refinements for inspection. Similar RNA-seq resources have been invaluable for pathogens such as *Salmonella* (Kröger et al., 2013). Robust annotations can then be linked to previously observed phenotypes or re-analyzed with existing datasets that are available for many pathogens, such as infection/stress-related gene expression/function datasets (Venturini et al., 2020).

## CONCLUSION

Our study represents a comprehensive blueprint for navigating the application, validation, and interpretation of translatome profiling methods for bacterial genome reannotation based on experimental data. We establish a new approach based on Ribo-seq to profile stop codons, which reveals ORFs that might be missed by *in silico* annotation as they are not apparent from the reference genome. We demonstrate that the *C. jejuni* genome encodes much more functional information than previously appreciated, for example in sORFs nested in longer genes, meaning much remains to be studied. We highlight the challenges in generating and utilizing robust genome annotations in the post-genomics world.

## METHODS

### Bacterial strains and culture conditions

All *Campylobacter jejuni* strains (**Table S7**) were routinely grown either on Müller-Hinton (MH) agar plates or with shaking at 140 rpm in *Brucella* broth (BB) at 37°C in a microaerobic atmosphere (10% CO_2_, 5% O_2_). All *Campylobacter* media was supplemented with 10 μg/ml vancomycin. Agar was also supplemented with marker-selective antibiotics [20 μg/ml chloramphenicol (Cm), 50 μg/ml kanamycin (Kan), 20 μg/ml gentamicin (Gm), or 250 μg/ml hygromycin B (Hyg)] where appropriate. *E. coli* strains (**Tables S7 & S8**) were grown aerobically at 37°C in Luria-Bertani (LB) broth or on LB agar supplemented with the appropriate antibiotics for marker selection.

### General recombinant DNA techniques and *C. jejuni* mutant construction

All plasmids generated and/or used in this study are listed in **Table S8**. Oligonucleotide primers (Sigma) are listed in **Table S9**. DNA constructs and mutations were confirmed by Sanger sequencing (Macrogen, Microsynth). Restriction enzymes, *Taq* polymerase for validation PCR, and T4 DNA ligase were purchased from NEB. For cloning purposes, Phusion high-fidelity DNA polymerase was used (Thermo Fisher Scientific). All *C. jejuni* mutant strains (deletion, chromosomal 3×FLAG-tagging, complementation by heterologous expression from the unrelated *rdxA* locus, **Table S7**) were constructed by double-crossover homologous recombination with DNA fragments introduced by electroporation into a *C. jejuni* strain NCTC11168 background. Details about *C. jejuni* mutant construction as well as transformation protocols are listed in the **Supplementary Methods**. For C-terminal epitope tagging, a 3×FLAG, SPA as used previously (Hemm et al., 2008; Zeghouf et al., 2004), or superfolder GFP (sfGFP, (Pédelacq et al., 2006)) sequence was fused to the penultimate codon of sORFs at their native locus by homologous recombination with an overlap PCR product. In some cases, SPA* was used, where the second codon of the SPA tag (ATG) was mutated to GCG.

### Total protein analysis by SDS-PAGE and western blotting

Bacterial cells were collected from cultures at the following densities (OD_600_): Lag: 0.1, eExp: 0.25, Exp: 0.5, Stat: 0.8 or from overnight cultures and resuspended in 1× protein loading buffer (62.5 mM Tris-HCl, pH 6.8, 100 mM DTT, 10% (v/v) glycerol, 2% (w/v) SDS, 0.01% (w/v) bromophenol blue). Samples corresponding to an OD_600_ of 0.1/0.2 were separated on 12% SDS-polyacrylamide (PAA) gels, or on 16% separating/4% stacking Tricine-SDS-PAGE gels without urea (Schägger, 2006) (stacking - 30V, separation 50-120V). Separated proteins were transferred to a nitrocellulose membrane by semidry blotting (Peqlab). After transfer, membranes were stained for 5 minutes in Ponceau S (Serva) to visualize transferred proteins (0.25% w/v Ponceau S, 5% acetic acid). Membranes were then blocked with 10% (w/v) milk powder in TBS-T (Tris-buffered saline-Tween-20) and incubated overnight with primary antibody in 3% BSA/TBS-T (monoclonal mouse anti-FLAG, 1:1,000; Sigma-Aldrich, #F1804-1MG, RRID:AB_262044 or monoclonal mouse anti-GFP, 1:1,000, Roche #11814460001, RRID:AB_390913) at 4°C. Washed membranes (TBS-T) were then incubated with secondary antibody (sheep polyclonal, anti-mouse IgG horseradish peroxidase (HRP) conjugate, 1:10,000 in 3% BSA/TBS-T; GE Healthcare, #RPN4201). Blots were developed using enhanced chemiluminescence reagent on a LAS-4000 Chemiluminescence Imager (GE Healthcare). An antibody specific for GroEL (rabbit polyclonal, 1:10,000 in 3% BSA/TBS-T; Sigma-Aldrich, #G6532-5ML, RRID:AB_259939) with an anti-rabbit IgG (goat polyclonal, 1:10,000 in 3% BSA/TBS-T; GE Healthcare, #RPN4301, RRID:AB_2650489) secondary antibody was used as a loading control. For size estimation, a Spectra™ Multicolor Low Range Protein Ladder or Prestained Protein Marker (Thermo Fisher Scientific) were loaded on Tricine-PAGE or SDS-PAGE gels, respectively. WB analysis of at least two independent biological replicates was performed and a protein was called as translated when it was detected in both replicates.

### Growth and cell harvest for ribosome profiling

For Ribo-seq, *C. jejuni* NCTC11168 WT or Δ*cmeB* mutant strains were grown in BB to log phase (OD_600_ ∼0.4-0.5). Full details are provided in the **Supplementary Methods**. For Ribo-seq with Cm, *C. jejuni* NCTC11168 WT cells were treated with 1 mg/ml Cm for 5 min at 37°C under microaerobic conditions, followed by immediate chilling for 10 min by mixture with an equal volume of crushed ice containing 1 mg/ml Cm. Cells were recovered by centrifugation and snap-frozen in liquid N_2_. For the TIS(Ret) experiment, Δ*cmeB* mutant cells were treated with 12.5 μg/ml Ret (Sigma CDS023386) for 10 min, followed by recovery by fast-filtration and snap freezing in liquid N_2_ as described previously (Gelhausen et al., 2022; Venturini et al., 2020). For the TIS/TTS experiment, *C. jejuni* NCTC11168 WT cultures were treated with peptide (NovoPro BioSciences, Shanghai; 50 μM final concentration, 10 min), immediately chilled in an ice bath with swirling for 3 minutes, recovered by centrifugation, and snap-frozen in liquid N_2_. MIC determination for Ret, Onc, and Api is described in the **Supplementary Methods**. Three independent biological replicates (cultures) were used for translatomics experiments.

### Processing of cell pellets and isolation of monosomes and footprints

Processing of cell pellets was performed generally as described previously (Gelhausen et al., 2022; Venturini et al., 2020). Cm was omitted from lysis buffers for TIS and TTS experiments, but used at 1 mM for Ribo-seq(Cm). PNP-GMP (guanosine 5′-[β,γ-imido]triphosphate, 3 mM, Sigma) was included in the lysis buffer of TIS(Ret) samples. Frozen cells were mixed with frozen lysis buffer (100 mM NH_4_Cl, 25 mM MgCl_2_, 20 mM Tris-HCl, pH 8, 0.1% NP-40, 0.4% Triton X-100) supplemented with 50 U DNase I (Thermo Fisher Scientific) and 500 U RNase inhibitor (moloX, Berlin). For Ribo-seq(Cm) and TIS(Ret), cells were lysed in a MM-400 metal ball mill (Retsch) for 5 rounds at 15 Hz for 3 min with chilling of the mill in liquid N_2_ between rounds. Lysates were thawed by incubation in a water bath at 30°C for 2 min and centrifuged at 10,000 *g* for 5 minutes. For the TIS(Onc)/TTS experiment, cells were lysed using a FastPrep system (MP Biomedical) with lysing Matrix B, speed 4, for 3 × 20s. Clarified lysates (approximately 15 A_260_ units) were digested with 800 U/A_260_ MNase (New England Biolabs) for 1.5 h (25°C, shaking at 14,500 rpm). Digests were stopped with EGTA (final concentration, 6 mM), immediately loaded onto 10-55% (w/v) sucrose density gradients freshly prepared in sucrose buffer (100 mM NH_4_Cl, 25 mM MgCl_2_, 5 mM CaCl_2_, 20 mM Tris-HCl, pH 8, 2 mM dithiothreitol), and centrifuged (35,000 rpm, 2.5 h, 4°C) in a Beckman Coulter Optima L-80 XP ultracentrifuge and SW40 Ti rotor. Gradients were fractionated (Gradient Station *ip*, Biocomp) and the 70S monosome fraction (identified by following fraction A_260_) was immediately frozen in liquid N_2_.

### RNA isolation and cDNA library preparation for translatomics

RNA was extracted from fractions or cell pellets for total RNA using hot phenol-chloroform-isoamyl alcohol or hot phenol, respectively, as described previously (Sharma et al., 2007; Vasquez et al., 2014). Total RNA was digested with DNase I, depleted of rRNA (Ribo-zero Bacteria, Illumina) and fragmented (Ambion 10× RNA Fragmentation Reagent) according to the manufacturer’s instructions. Monosome RNA and fragmented total RNA was size-selected (26-34 nt) on 15% polyacrylamide/7M urea gels as described previously (Ingolia et al., 2012). Libraries were prepared by vertis Biotechnologie AG (Freising, Germany) using a small RNA protocol (Ribo-seq(Cm) or TIS(Ret) experiment) or an Adaptor Ligation protocol (TIS(Onc)/TTS experiment) and sequenced on a NextSeq500 instrument (high-output, 75 cycles) at the Core Unit SysMed at the University of Würzburg.

### General processing of ribosome profiling data

Sequencing data was processed and analyzed with HRIBO (version 1.4.4) (Gelhausen et al., 2021), an analysis workflow for bacteria based on snakemake (RRID:SCR_003475) (Köster and Rahmann, 2018). All required dependencies are automatically downloaded from *bioconda* (RRID:SCR_018316) (Grüning et al., 2018) and docker (Merkel, 2014). Adapter trimming was performed with cutadapt (version 2.1, RRID:SCR_011841) (Martin, 2011) and reads were mapped with *segemehl* (version 0.3.4, RRID:SCR_005494) (Hoffmann et al., 2009). Multi-mapping reads and those mapping to ribosomal RNA were removed using SAMtools (version 1.9, RRID:SCR_002105) (Li et al., 2009). The reference genome sequence NC_002163.1 (ASM908v1) (Gundogdu et al., 2007; Parkhill et al., 2000) was used in combination with a custom annotation file including previously-annotated 5’UTRs and sRNAs (Dugar et al., 2013) for Ribo-seq data analysis. To generate this file, the NCBI annotation (2014-03-20) was combined with a 5’UTR and sRNA annotation generated based on dRNA-seq data (Dugar et al., 2013). In brief, primary transcriptional start sites in the NCTC11168 genome as well as the strand information (Table S4) was used to calculate 5’UTR end positions based on UTR lengths to identify 5’UTR regions. Resulting 5’UTR regions were exported in GFF3 format. To generate the sRNA annotation file, the annotation table for strain NCTC11168 from (Dugar et al., 2013) (Table S11) was used. CJas_Cj0168c, not detected by NB, was excluded from the final sRNA annotation file. The three .gff files were combined and attribute columns were updated to fit the GFF3 standard. Furthermore, four housekeeping RNAs (SRP, RnpB, tmRNA, and TPP riboswitch) were added to the custom annotation (also from Table S11, (Dugar et al., 2013)). Comparison of the NCBI genome annotation used for analysis (2014-03-20) to the one currently available at NCBI (updated 2021-09-11, downloaded 2022-11-07) revealed that they are identical except for two gene fusions that were replaced by single pseudogenes (*uxaA’* (Cj0482)/*uxaA* (Cj0483), now Cj0483 pseudogene and *metC’* (Cj1392)/*metC* (Cj1393), now Cj1392 pseudogene). Summary statistics can be found in **Table S10**.

### Ribo-seq-based ORF prediction

ORFs were called from Ribo-seq libraries using two prediction tools included in HRIBO: DeepRibo (Clauwaert et al., 2019) and Reparation_blast, an adapted version of REPARATION (Ndah et al., 2017), which uses blast instead of usearch (https://github.com/RickGelhausen/REPARATION_blast) (Gelhausen et al., 2022). Predictions were generated from normal Ribo-seq data from all datasets ((Ribo-seq (Cm), control (no drug) libraries from the TIS/TTS experiments). Three replicates were used for Ribo-seq(Cm) and TIS(Ret) experiment data, while only the first available replicate for the TIS(Onc)/TTS experiment was used to generate predictions. The results of both DeepRibo and REPARATION from all analyzed experiments/replicates were aggregated into Excel tables with additional information for each detected ORF, *e*.*g*., expression, translational efficiency, DeepRibo score for subsequent filtering (see section on Filtering below).

### Detection of TIS/TTS sites and associated ORFs

For detection of TIS/TTS, we adapted previous peak detection methods (Gelsinger et al., 2020; Weaver et al., 2019). All programming scripts are available at https://github.com/RickGelhausen/StartStopFinder. Briefly, we used single-nucleotide mapping coverage files generated by HRIBO (version 1.4.4) (Gelhausen et al., 2021) for read 3’ends and read lengths providing the sharpest enrichment near the expected offset positions with respect to the start/stop codon within each experiment, based on metagene analysis (Becker et al., 2013). Selected mapping files and offsets were then used for peak detection at start codons (ATG/GTG/TTG) at intervals of 5 nt. For each interval with a non-zero peak height, the next in-frame stop codon was identified to generate a corresponding ORF (of any length). For TTS-based sORF detection, a similar approach was taken (TGA/TAA/TAG codons), except potential sORFs with a stop codon within 25 nt of an annotated stop codon were omitted to exclude predictions arising from read-through and stop codons with longest potential length ≤71 codons or shortest potential length ≥11 codons were investigated manually for start codon peaks, due to ambiguity in assigning start codons. For details, including a list of read-lengths and offsets for all experiments, see the **Supplementary Methods**.

### Filtering of Ribo-seq/TIS/TTS ORF predictions

To generate a manageable list of candidate sORFs (11-71 codons) from HRIBO Ribo-seq-based predictions, we applied the following expression cutoffs (mean TE and RNA-seq RPKM (within an experiment) ≥1 and ≥30, respectively, see also **Figs. S8A & S8D**). Candidates were required to be detected by REPARATION or by DeepRibo (with a Score >0) in at least one replicate of the experiment. We also required candidates to be detected in a Ribo-seq library for at least two out of the three experiments (Ribo-seq(Cm), TIS(Ret), and TIS(Onc)/TTS). To generate a list of sORF candidates based on TIS for curation, we applied the following cutoffs (see also **Figs. S8B & S8D**): ORF length between 11-71 codons, peak height (Ret or Onc TIS library) ≥20, a peak in the TIS library only or a log2FC (fold-change) ≥1 in at least two replicates (TIS(Ret) experiment) or the single TIS/TTS(Onc/Api) replicate used for predictions. For TIS predictions, we also manually inspected the top 10 candidates (detected in all three replicates with a TIS peak only or log2FC>1, sorted by TIS peak height) shorter than the DeepRibo/REPARATION length cutoff of 11 codons (4-10 codons). To filter TTS sites, we applied the following criteria (see also **Fig. S8D**). Because of uncertainty in assigning start codons, predictions were associated with a longest and a shortest potential ORF. Candidate sORFs were filtered for those with the longest potential ORF ≤71 codons and the shortest potential ORF ≥11 codons. The minimum TTS peak height was required to be ≥5 reads. For TTS(Mono), all candidates with a peak at the same position in the corresponding TIS(Onc) library and all internal out-of-frame candidates were excluded.

### Manual curation of Ribo-seq/TIS/TTS-based sORF predictions

The 421 filtered sORF predictions were inspected based on coverage files for one replicate of each experiment and were inspected using criteria previously used for bacterial Ribo-seq data (Gelhausen et al., 2022). Coverage files were loaded together in IGB (Integrated Genome Browser (Freese et al., 2016), https://www.bioviz.org/) along with ORF, sRNA, and TSS annotations (Dugar et al., 2013; Gundogdu et al., 2007). ORFs were called as ‘translated’ using the following criteria. First, the Ribo-seq signal was generally required to be comparable to or higher than the transcriptome library (*i*.*e*., TE approx. 1). Second, the shape of the Ribo-seq coverage over the ORF was considered: ORFs with Ribo-seq coverage near the start codon and/or restricted within ORF boundaries (and excluded from 5′/3′UTRs) were prioritized, even if the TE was <1. For TIS peaks, the offset with respect to the start codon (+/-2 nt around the ideal offset based on metagene analysis) as well as peak height and enrichment (vs. the Ribo-seq library) were evaluated. In addition, we evaluated the enrichment of TIS peaks with respect to surrounding noise. For TTS peaks, the offset with respect to the stop codon, +/-2 nt based on metagene analysis, as well as peak height and presence of a TIS(Onc) peak were investigated. We also evaluated local enrichment compared to the surrounding noise, as well as whether the peak might reflect stop codon read through. TTS peaks with a TIS(Onc) peak at the same position were also discarded. For predictions derived from TTS data only, the corresponding best start codon was identified by inspection of Ribo-seq and TIS coverage. For all sORF predictions, the presence of a potential RBS and TSS position were also considered.

### Additional bioinformatics and analysis

RBS motifs were predicted using MEME (version 5.4.1, RRID:SCR_001783) (Bailey et al., 2009). Strains and genome accessions used for conservation analysis are listed in **Table S6**. Amino acid sequences for *C. jejuni* NCTC11168 annotated (Gundogdu et al., 2007) or novel (**Table S1**) small proteins were used as a query for tBLASTn searches at NCBI (https://blast.ncbi.nlm.nih.gov/Blast.cgi; RRID:SCR_011822). Default parameters were used, except for the following modifications, based on those previously used for bacterial sORFs (Allen et al., 2014): Expect threshold (E-value) ≤ 100, word size 3, no coverage cutoff, filter for low complexity off). Annotated homologs were detected by BLASTp at NCBI (RRID:SCR_001010) using default parameters. For a “yes” match, both an identical same length and identical amino acid sequence was required. For a partial match, an E-value <100 was still required, but the length requirement was omitted (*i*.*e*., the matching ORF could be the longer parental/overlapping ORF for an internal in-frame candidate). Subcellular localization of small proteins was predicted using pSORTb v3.0 (RRID:SCR_007038) with default parameters for Gram-negative bacteria (Yu et al., 2010).

### Mass spectrometry-based proteomics

Small proteins were identified in *C. jejuni* soluble protein extracts, generated by lysis using a FastPrep Homogenizer (MP-Biomedicals), by MS from bacteria growing in log phase in BB. Two different techniques for pre-fractionation of proteins were applied: (i) separation of soluble proteins by one dimensional (1D) SDS-PAGE and in-gel digestion with trypsin or chymotrypsin (see “gel-based approach” described in (Fuchs et al., 2021), or (ii) fractionation of proteins on a GELFREE 8100 fractionator (Expedeon) with 10% Tris acetate cartridges and trypsin digestion. Digestion of proteins using a GELFREE 8100 fractionator was performed in protein low binding tubes (Eppendorf, Hamburg, Germany) using the Single-Pot Solid-Phase-enhanced Sample Preparation technique described previously (Hughes et al., 2014) with modifications as described in the **Supplementary Methods**. Peptide fractions were analyzed using the Orbitrap Fusion MS coupled to a Dionex Ultimate 3000 nHPLC system (Thermo Fisher Scientific Inc., Waltham, Massachusetts, USA) as described previously (Fuchs et al., 2021) with modifications as described in the **Supplementary Methods**.

For identification of small proteins based on MS/MS data, we used the fully automated bacterial proteogenomics workflow SALT & Pepper (https://gitlab.com/s.fuchs/pepper; (Fuchs et al., 2021)), which includes protein database generation, database searching, peptide-to-genome mapping, and result interpretation. MS- and MS/MS-data of all samples were searched by MaxQuant (Max Planck Institute of Biochemistry, Martinsried, Germany, www.maxquant.org, version 1.5.2.8, RRID:SCR_014485) against a database with *C. jejuni* annotated protein sequences from NCBI (downloaded on 01-09-2020) and sORFs that were predicted using our Ribo-seq data and a translational database (TRDB) of the full coding potential of the *C. jejuni* genome generated by six-frame translation from stop codon to stop codon with a minimum length of 9 aa generated by SALT (https://gitlab.com/s.fuchs/pepper). Three independent biological replicates were used for analysis. Full details of MS-based proteomics can be found in the **Supplementary Methods**.

### Microscopy

For FITC (fluorescein isothiocyanate) labeling of *C. jejuni*, ∼1.5 - 2 × 10_8_ cells, grown in log phase in BB + 10 µg/ml vancomycin, were harvested by centrifugation (6600 *g*, 5 minutes at room temperature) and washed once with 1× PBS. Freshly prepared 10 mg/ml FITC (Sigma) in 100% ethanol was diluted to 0.1 mg/ml in 1× PBS. Bacteria were resuspended in this solution for 30 min (37°C, microaerophilic conditions, shaking), washed twice with 1× PBS, and fixed with 4% paraformaldehyde for 1 hour at room temperature in the dark. After washing once with 1× PBS, cells were resuspended in 1× PBS and placed on an agarose pad (1% agarose in 1× PBS) and imaged with a laser scanning Leica TCS SP5 II confocal microscope (Leica Microsystems).

### Motility assays

Liquid cultures of each strain were grown to log phase in BB media + 10 µg/ml vancomycin. Next, 1 μl of bacterial culture was inoculated into a soft-agar plate (BB broth + 0.4% Difco agar, 10µg/ml vancomycin). Plates were incubated right-side-up at 37°C under microaerobic conditions until halo formation could be observed (approximately 24 hrs post-inoculation). Each halo radius of technical triplicates was measured two times and averaged to give the mean swimming distance per strain. Motility assays were performed in three independent biological replicates. Student’s t-test (unpaired, two-tailed) was used to assess significance.

### Co-immunoprecipitation (coIP) for investigation of protein-protein interactions

Lysates were prepared from *C. jejuni* strains, grown to log phase, carrying chromosomally epitope-tagged versions of CioA and/or CioY (CioA-SPA & CioY-sfGFP and reciprocal version CioY-SPA & CioA-sfGFP), with a FastPrep system (MP Biomedical, matrix B and lysis buffer with 1% DDM (n-dodecyl-B-D-maltoside). Lysates of the untagged wild-type strain as well as the corresponding sfGFP-only tagged strains (CioA-sfGFP or CioY-sfGFP alone) were used as a control for unspecific binding. SPA-tagged protein was pulled down from clarified lysates with an anti-FLAG antibody (Sigma-Aldrich, #F1804-1MG, RRID:AB_262044) bound to Protein A-Sepharose beads (Sigma-Aldrich, #P6649). Co-purification was investigated by western blot with an anti-GFP antibody (Roche #11814460001, RRID:AB_390913). Two independent biological replicates were performed. Full details can be found in the **Supplementary Methods**.

### Protein complex and structure prediction with AlphaFold2

Structural predictions of protein complexes were performed at ColabFold (https://colab.research.google.com/github/sokrypton/ColabFold/blob/main/AlphaFold2.ipynb#scrollTo=KK7X9T44pWb7, accessed 2022-03-26) using AlphaFold2 and AlphaFold2-multimer (Mirdita et al., 2022). Standard settings were used (msa_mode: MMseqs2 (UniRef+Environmental); pair_mode: unpaired+paired; model_type: auto; num_recycles: 3). The best ranked structure prediction was selected, where pLDDT values for all proteins in the complex were evaluated in addition. Graphics of structural predictions, structures from PDB (6RKO), as well as overlays were generated in Pymol (RRID:SCR_000305).

## ACKNOWLEDGEMENTS

We thank Gaurav Dugar, Sandy Pernitzsch, and Lydia Hadjeras for discussions on Ribo-seq protocols, as well as Thorsten Bischler and Diego Gelsinger for guidance on Ribo-seq and TIS data analysis. We thank Daniel Wilson for discussions regarding TTS profiling and Julian Langner for fruitful discussions regarding oxidase complexes as well as for providing reagents. We are also grateful to Susan Gottesman for insightful comments on the manuscript.

## AVAILABILITY OF DATA AND MATERIALS

Ribo-seq data have been deposited at the NCBI Gene expression Omnibus (GEO) under the Accession GSE208756. The mass spectrometry proteomics data have been deposited to the ProteomeXchange Consortium (http://proteomecentral.proteomexchange.org) via the PRIDE partner repository (https://www.ebi.ac.uk/pride/archive) (Vizcaíno et al., 2016) with the dataset identifier PXD036790. Differential RNA-seq data (Dugar et al., 2013) was recovered from GEO (GSE38883). All programming scripts are available at https://github.com/RickGelhausen/StartStopFinder.

## COMPETING INTERESTS

The authors declare no competing interests.

## AUTHOR CONTRIBUTIONS

K.F., S.L.S., and C.M.S. designed the research. K.F., S.L.S, E.F., P.K., and A.K. performed lab work. K.F., S.L.S., R.G., A.K., M.K., S.F., and F.E. analyzed data. K.F., S.L.S., and C.M.S. interpreted the data and wrote the manuscript, which all co-authors revised. R.B., S.E., and C.M.S. supervised research and provided resources.

## FUNDING INFORMATION

Small proteins and Ribo-seq research in the Cynthia M. Sharma laboratory is supported by an individual project grant within the Deutsche Forschungsgemeinschaft (DFG) priority program SPP2002 “Small proteins in prokaryotes, an unexplored world” to C.M.S. (SH580/8-1 and SH580/8-2). This work was also supported by the Z2 Central Project “Ribosome Profiling and Bioinformatics’’ within the SPP2002 (awarded to C.M.S. SH580/7-1 and 7-2 and R.B. BA2168/21-2). K.F. was supported by a grant of the German Excellence Initiative to the Graduate School of Life Sciences, University of Würzburg. Computational resources were provided by the BMBF-funded de.NBI Cloud within the German Network for Bioinformatics Infrastructure (de.NBI) (031A532B, 031A533A, 031A533B, 031A534A, 031A535A, 031A537A, 031A537B, 031A537C, 031A537D, 031A538A) to R. B. MS-based small protein analyses were supported by an individual project grant within the DFG priority program SPP2002 “Small proteins in prokaryotes, an unexplored world” to S.E. and S.F. (EN 712/4-1), by a grant of the GRK PROCOMPAS (DFG) to S.E., and an institutional grant (INST 188/365-1 FUGG DFG) to S.E.

## Notes

### Competing Interest Statement

The authors have declared no competing interest.

