## Supplementary Materials for "Complementary Ribo-seq approaches map the translatome and provide a small protein census in the foodborne pathogen *Campylobacter jejuni*"

**This document contains:**

**Supplementary Figures S1-S12**

**Supplementary Methods**

**Supplementary References**

### Supplementary Figures

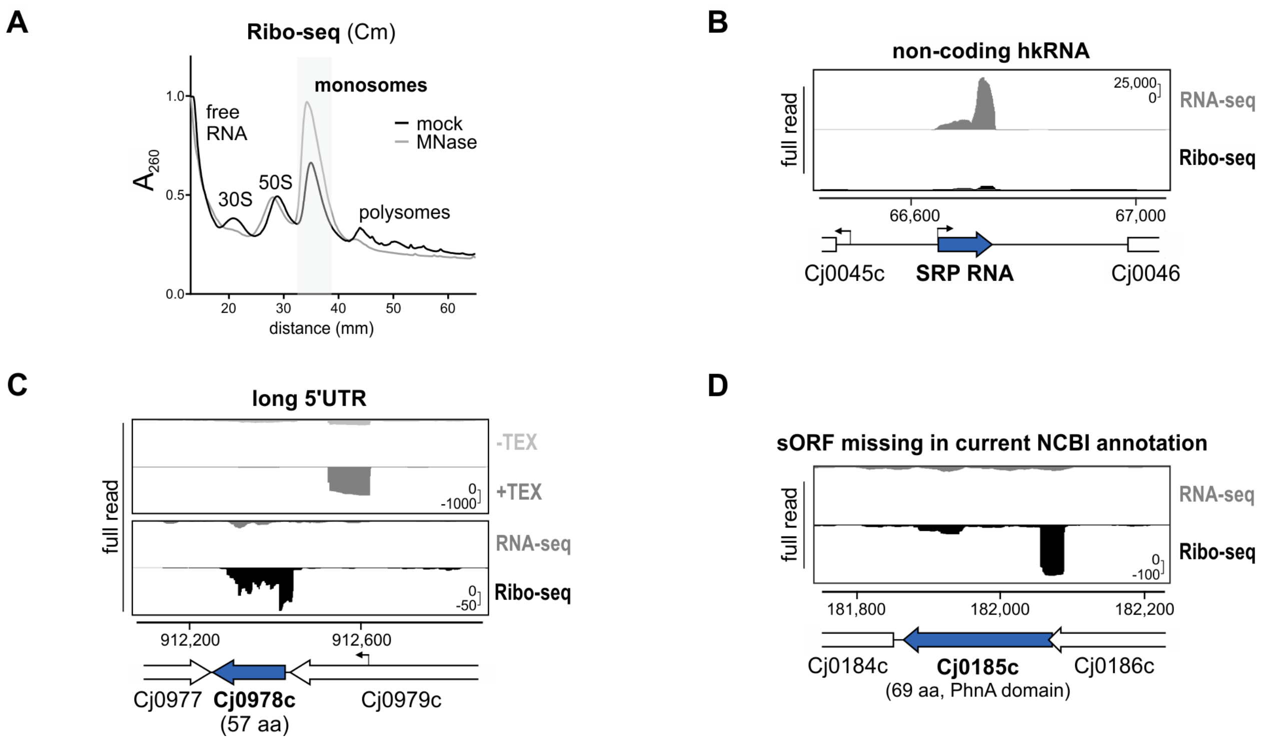


**Supplementary Figure S1. A *C. jejuni* Ribo-seq protocol captures translating ribosomes and distinguishes coding from non-coding features. (A)** Polysome profiles for mock- and micrococcal nuclease (MNase)-treated lysates from *C. jejuni* strain NCTC11168 WT. Cells were harvested from log phase cultures in rich medium. Lysates were separated on 10-55% sucrose density gradients by ultracentrifugation and the RNA content of fractions was measured ($A_{260}$) to identify complexes. Representative of three independent experiments. **(B)** cDNA reads for the non-coding housekeeping RNA (hkRNA) component of the signal recognition particle (SRP) are mainly restricted to RNA-seq, rather than Ribo-seq, libraries. Representative of three independent experiments. **(C)** MNase digestion of unprotected RNA validates a long (~200 nt (Dugar et al., 2013)) 5′UTR of sORF Cj0978c (57 aa). **(D)** Ribo-seq suggests that Cj0185c, absent from the current *C. jejuni* NCTC11168 NCBI annotation, is translated from an operon with Cj0184c and Cj0186c. For all screenshots, y-axis scales represent rpm (reads per million) and coverage is representative of three independent experiments.

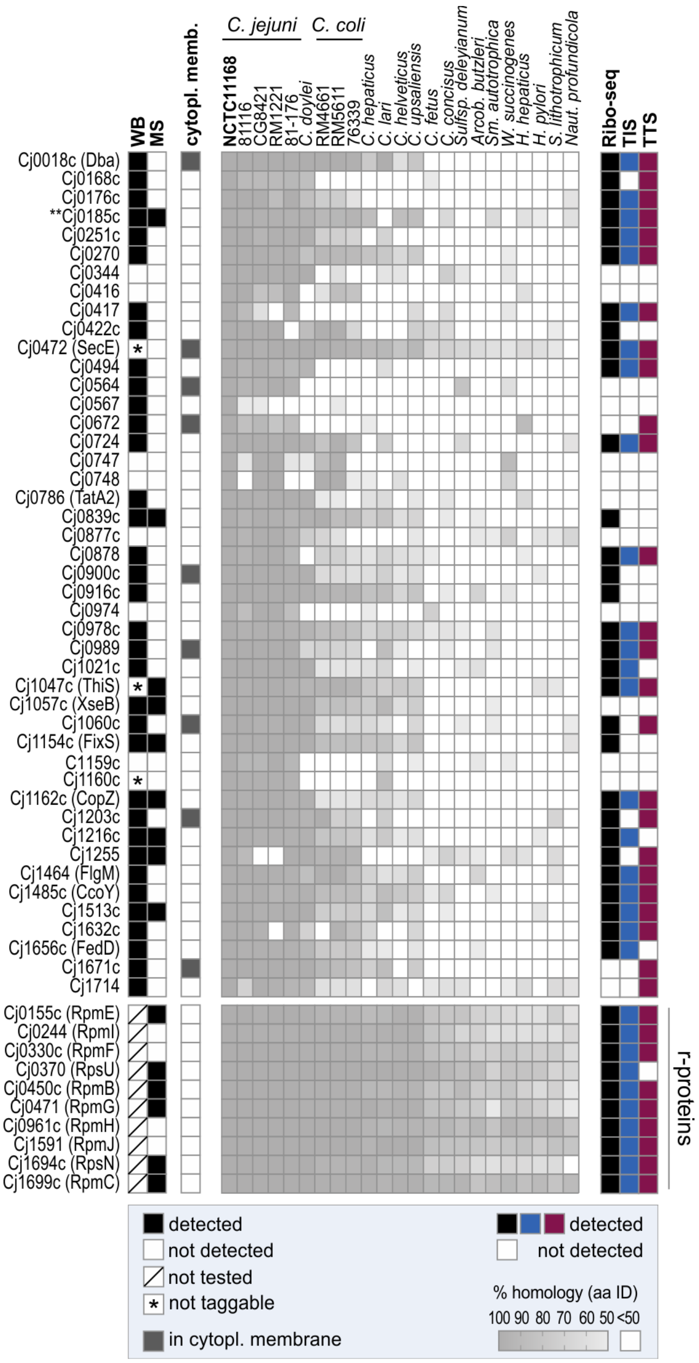


**Supplementary Figure S2. Overview of results obtained from the different sORF detection approaches, additional features, and conservation for annotated CJsORFs.** WB: western blot (detected at any growth phase tested). MS: mass spectrometry detection in log phase. For conservation, sequences from *C. jejuni* NCTC11168 were used for tBLASTn against the indicated strains (see also **Table S6**). tBLASTn: word size 3, E-value ≤ 100. Cytoplasmic membrane: PSORTb predictions for subcellular localization (Yu et al., 2010). Cj0185c, absent from current NCBI annotation, is included (**). Genes for which no clones for C-terminal 3xFLAG/SPA fusions were obtained after several transformation trials, are indicated as not taggable (*).

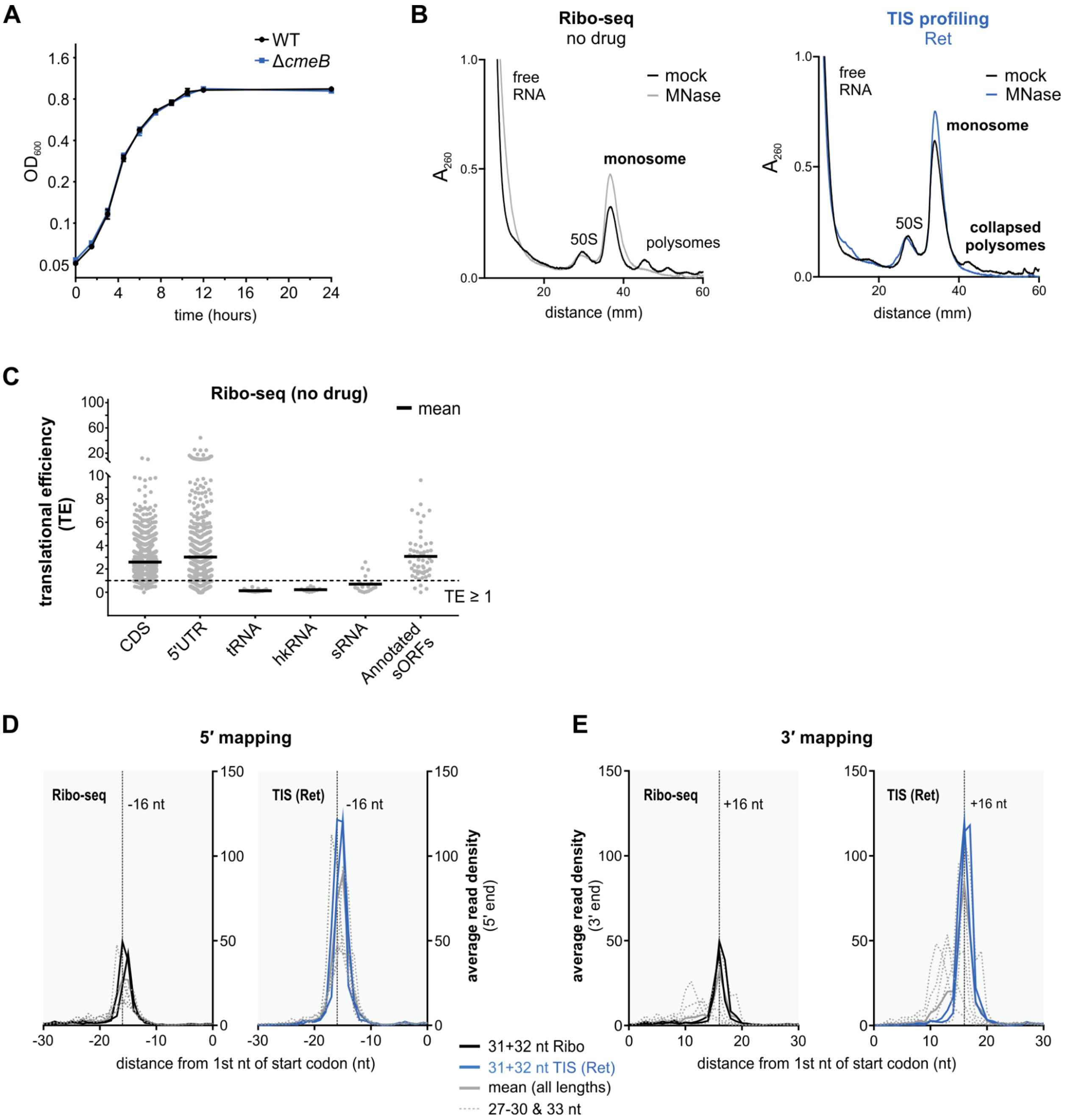


**Supplementary Figure S3. Establishing Ret-mediated TIS profiling in *C. jejuni*. (A)** Growth of a *C. jejuni* Δ*cmeB* mutant vs. the parental WT strain under standard conditions in rich media. Ribo-seq cultures were harvested at $OD_{600}$ ~0.4. Error bars indicate the standard deviation of three independent cultures. **(B)** Polysome profiles (-/+ MNase treatment) for untreated (no drug; *left*) or Ret-treated (12.5 µg/ml, 10 min; *right*) *C. jejuni* Δ*cmeB*. Lysates were separated on a 10-55% sucrose density gradient by ultracentrifugation and the RNA content of fractions was measured ($A_{260}$) to identify complexes. **(C)** Translational efficiency (TE: Ribo-seq/RNA-seq) for feature classes in the *C. jejuni* annotation (Dugar et al., 2013; Gundogdu et al., 2007) for one replicate of the no-drug control (no Cm) of the TIS(Ret) experiment. CDS: coding sequence. hk: housekeeping. **(D & E)** Metagene analysis of ribosome occupancy (no drug vs. Ret) at start codons for indicated read lengths (5′ read end mapping, panel **D**; 3′, panel **E**). Related to main **Fig. 2A**. Solid black: 31 or 32 nt-long reads for Ribo-seq (no drug). Solid blue: 31 or 32 nt reads for TIS profiling (Ret). Solid grey: mean of all lengths. Dashed grey lines: all other lengths (27-30 and 33 nt).

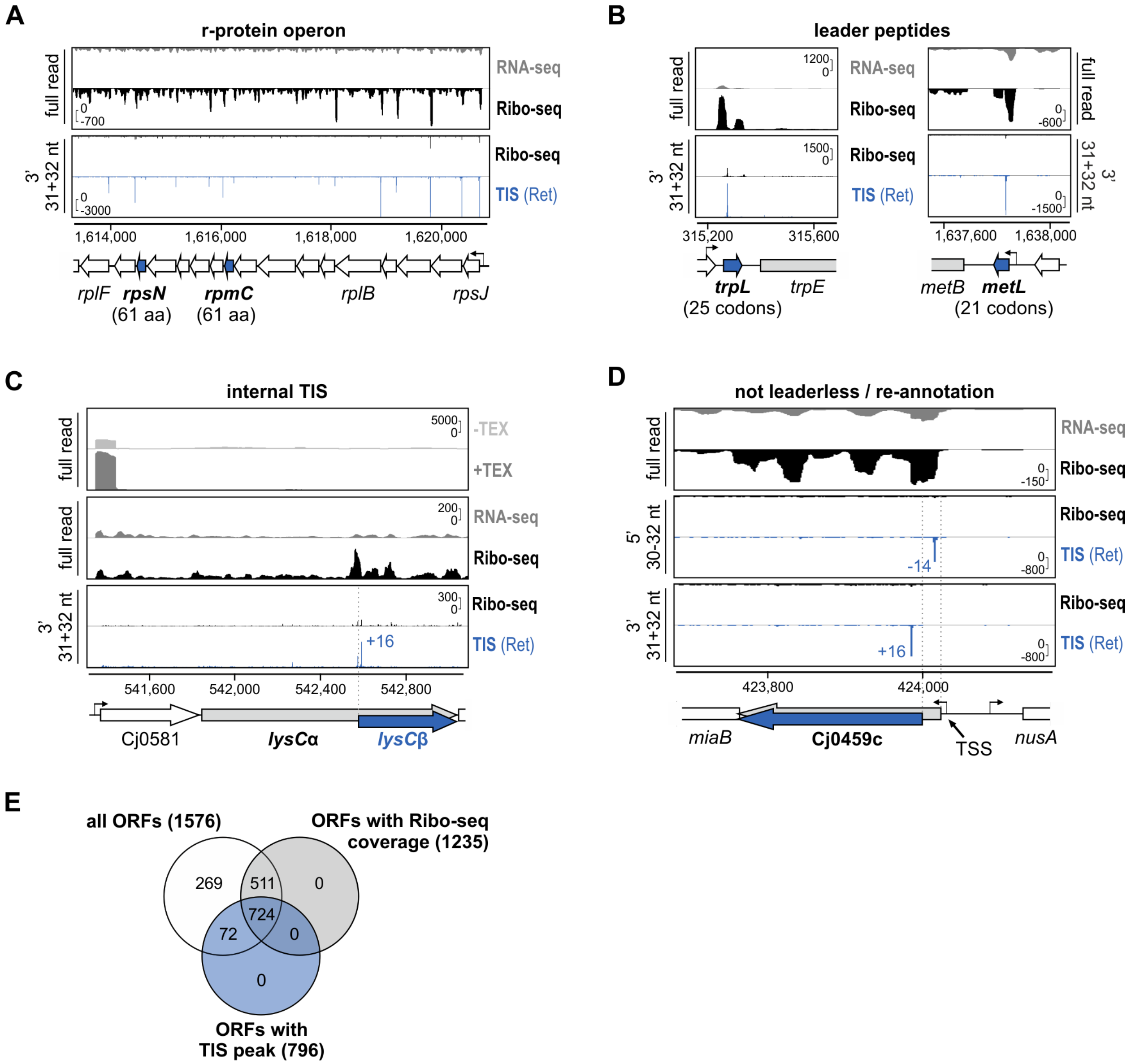


**Supplementary Figure S4. Ret-mediated TIS profiling reveals *C. jejuni* start codons. (A)** Ribosome occupancy across an r-protein operon. Two sORFs (*rpsN* & *rpmC*, both 61 aa) are highlighted. Top two tracks: full read coverage for paired RNA-seq and conventional Ribo-seq libraries. Bottom two tracks: 3'-end coverage for 31 + 32 nt reads for Ribo-seq and TIS libraries. Enriched peaks in TIS (Ret) vs. Ribo-seq (arrows) suggest translation initiation sites. **(B)** TIS detection of two previously predicted *C. jejuni* leader peptides/uORFs in amino acid biosynthesis operons (TrpL, *left;* MetL, *right*) (Porcelli et al., 2013). Related to main **Fig. 2B**. **(C)** TIS data reveals a potential 156 aa LysCβ isoform encoded in *lysC* (LysCα, Cj0582, 401 aa) generated via internal translation initiation as reported in other bacteria (Meydan et al., 2018). +16 nt: TIS peak. The *lysCβ* ORF (blue) was also predicted from Ribo-seq data. **(D)** TIS profiling suggests the predicted (Porcelli et al., 2013) leaderless gene Cj0459c (5′UTR: 6 nt) should be re-annotated as leadered with a 30 nt 5′UTR. Both 5′ and 3′ end coverage was used. For all screenshots, bent arrows: TSS based on dRNA-seq (Dugar et al., 2013). Y-axis scales represent rpm (reads per million). **(E)** Overlap of all annotated ORFs with ORFs with Ribo-seq coverage (TE ≥1; total RNA RPKM ≥30) and detected TIS(Ret) peaks.

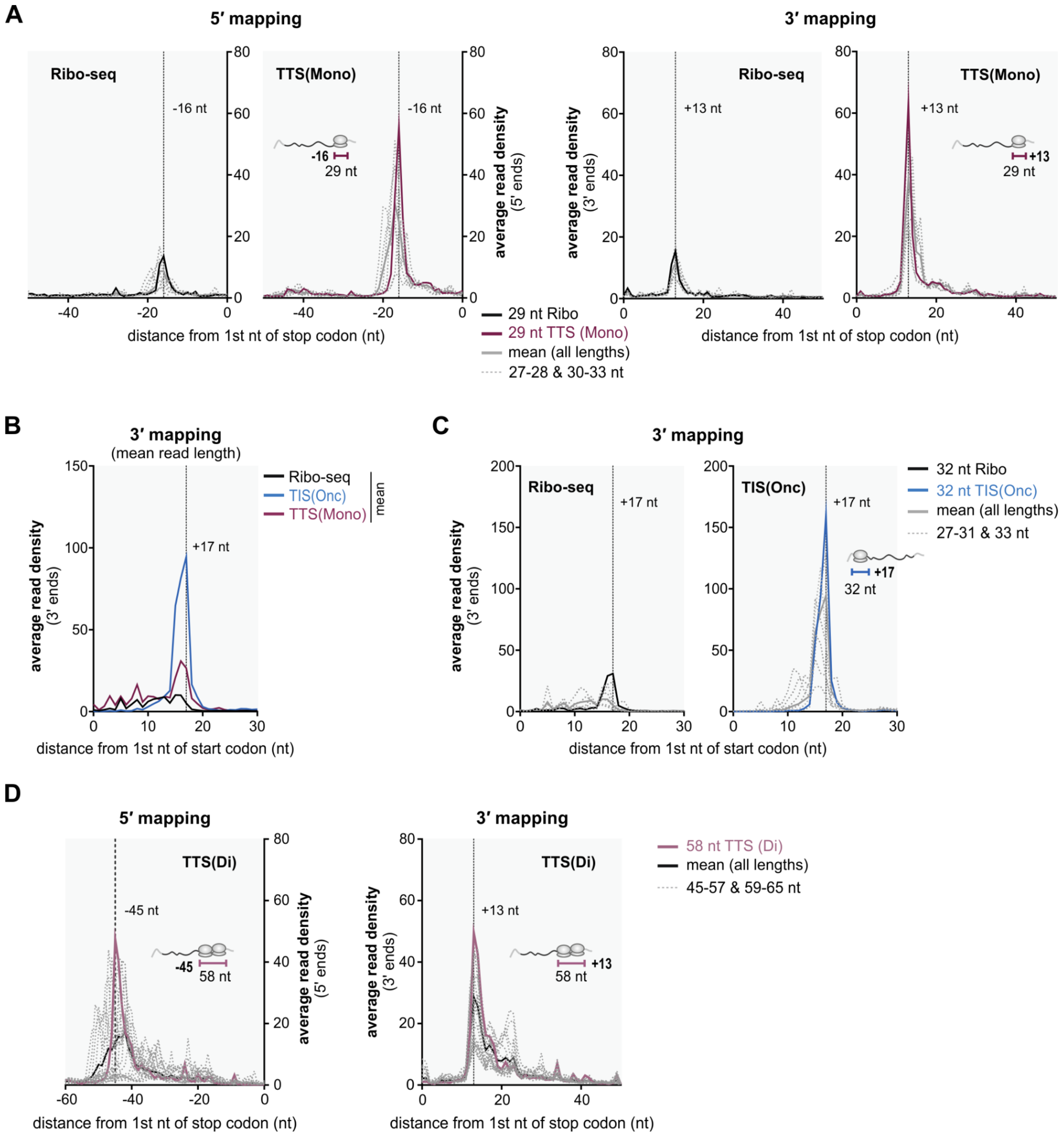


**Supplementary Figure S5. Metagene analysis of ribosome occupancy for offset and read length determination for the TTS profiling experiment. (A)** Ribosome occupancy near stop codons using 5′/3′ read end coverage for Ribo-seq vs. TTS monosome libraries (TTS(Mono)). Solid black lines: 29 nt reads (Ribo-seq). Solid red lines: 29 nt reads (TTS(Mono)). Solid grey lines: mean of all read lengths. Dashed grey lines: all other read lengths (27-28 and 30-33 nt). Related to main **Fig. 3C**. **(B)** Ribosome occupancy near start codons using 3′ read end coverage (mean of all read lengths) for the Ribo-seq, TIS(Onc), and TTS(Mono) libraries. **(C)** Ribosome occupancy near start codons using 3′ read end coverage for different read lengths in Ribo-seq vs. TIS(Onc) libraries. Solid black line: 32 nt reads of Ribo-seq. Solid blue line: 32 nt reads of TIS(Onc). Solid grey lines: mean of all read lengths. Dashed grey lines: all other read lengths (27-31 and 33 nt). **(D)** Ribosome occupancy near stop codons for different read lengths in the TTS disome library (TTS(Di)). Solid black lines: mean of all lengths. Solid red lines: 58 nt reads of TTS(Di). Dashed grey lines: all other read lengths (45-57 and 59-65 nt). Related to main **Fig. 4D**. Representative of three independent replicates.

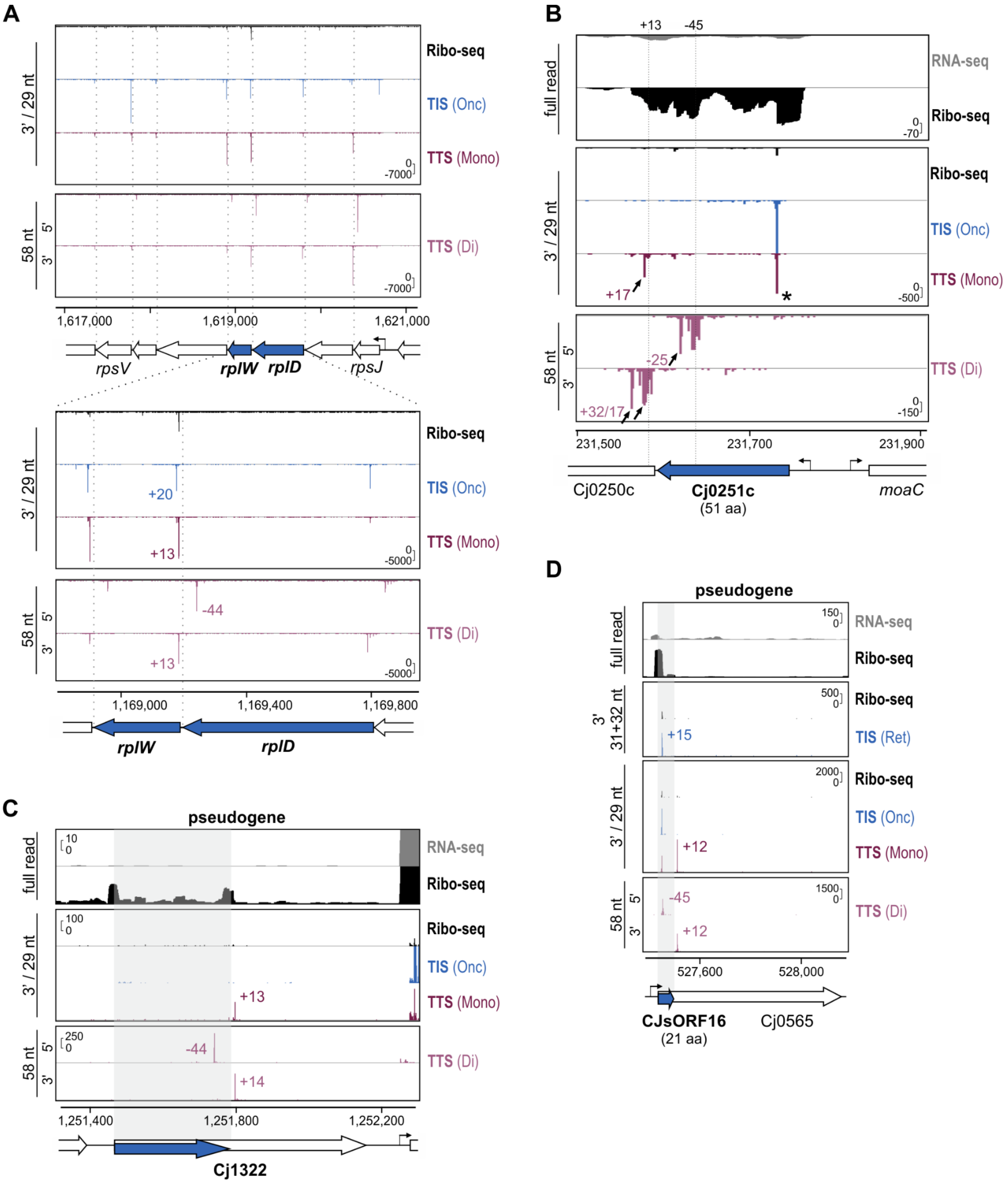


**Supplementary Figure S6. Api-mediated TTS profiling reveals *C. jejuni* stop codons. (A)** Single-nt (read 5′ or 3′ end) mapping coverage from the TTS experiment for an r-protein operon. Bottom: zoom-in on *rplD* ORF boundaries shows peaks at the expected offsets: +13 nt for TTS(Mono) and TTS(Di) (3′ end mapping); -44 nt for TTS(Di) (5′ end mapping). The indicated TIS peak at +20 nt (vs. the *rplD* stop codon) is at the expected +17 position for the *rplW* start codon. Dashed lines: stop codon positions. **(B)** TTS coverage for the Cj0251c sORF (51 aa) demonstrates stop codon readthrough induced by Api (arrows). Asterisk: partial enrichment at the start codon by Api. Dashed lines: expected TTS positions (+13 nt (monosomes and disomes, 3′ end) and -45 nt (disome 5′ ends)). **(C)** Translation of the Cj1322 pseudogene until an in-frame stop codon mutation in the reference genome. **(D)** TIS/TTS-based detection of a novel sORF (CJsORF16, 21 aa) generated from translation of a pseudogene (Cj0565) until an in-frame stop codon generated by mutation. For all screenshots, bent arrows: TSS based on dRNA-seq (Dugar et al., 2013). Y-axis scales represent rpm (reads per million).

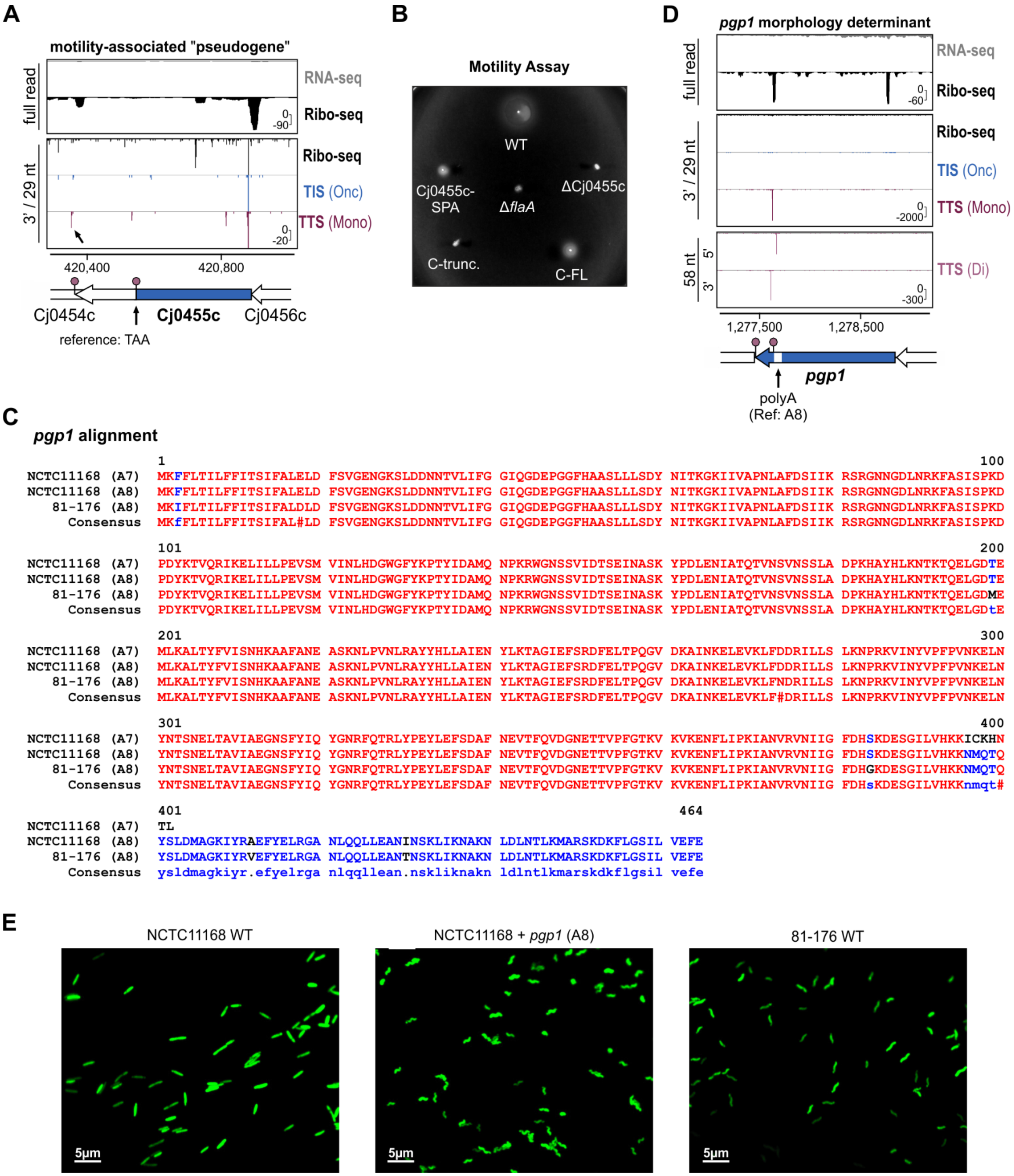


**Supplementary Figure S7. Translation status of motility and shape-determining proteins. (A)** TTS profiling supports full-length translation, beyond an annotated premature stop codon, of motility-related "pseudogene" Cj0455c. TTS(Mono) peak (arrow) associated with full-length Cj0455c. TAA: premature stop codon. Y-axis scales represent rpm (reads per million) (same for panel D). **(B)** Representative motility assay of a *C. jejuni* ΔCj0455c mutant complemented in the chromosome either with the full-length (C-FL; CAA) or the truncated (C-trunc.; TAA) Cj0455c isoform at unrelated *rdxA*. The C-terminal SPA-tagged Cj0455c strain is included. Δ*flaA*: non-motile control. Related to main **Fig. 4B**. Representative of three independent experiments. **(C)** Alignment of *pgp1* aa sequences from *C. jejuni* NCTC11168 (A8 reference and actual A7 determined by Sanger sequencing) and 81-176 using MultAlin (Corpet, 1988). Red: 100% aa identity. Blue: >50% aa identity. **(D)** TTS peaks are consistent with rod shape and *pgp1* genotype (*pgp1*-ON/OFF(A8/A7)) of the NCTC11168 WT isolate used for Ribo-seq/TTS. **(E)** Confocal micrographs for analysis of cell morphology of *C. jejuni* strains expressing *pgp1* alleles. Bacteria were stained with FITC (fluorescein isothiocyanate). Related to main **Fig. 4D**. NCTC11168 WT: straight isolate used in this study. NCTC11168 + *pgp1* (A8): WT carrying, in addition to the *pgp1* A7 allele at the native locus, a copy of *pgp1*-A8 from 81-176 at unrelated *rdxA*. 81-176 WT: WT (spiral) isolate.

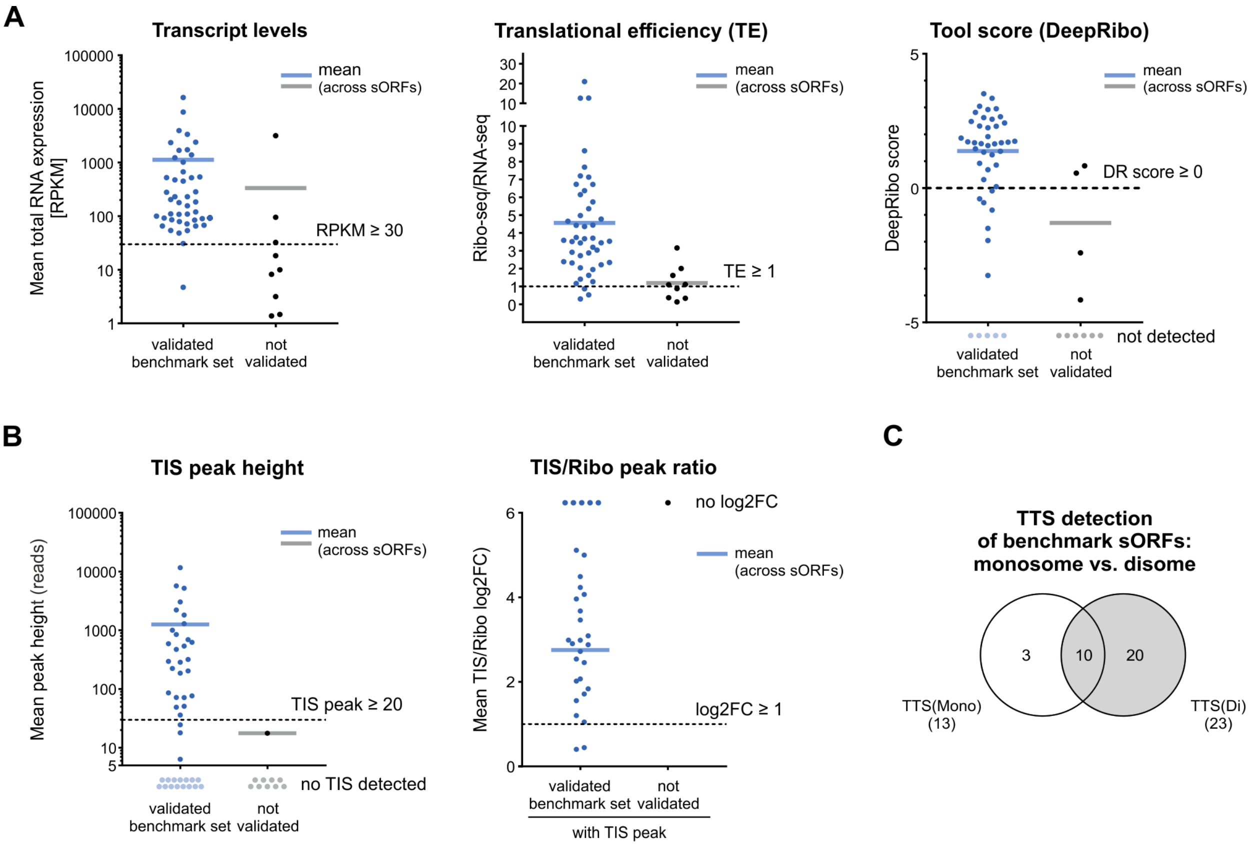


D

| | **Ribo-seq** Cm / no drug | **TIS profiling** Ret / Onc | **TTS profiling** Api (Mono / Di) |
|---|---|---|---|
| **Filtering criteria** | • ≤ 71 codons<br>• avg. TE ≥ 1, avg. RNA rpkm ≥ 30<br>• predicted in Reperation<br>• or DeepRibo (DR score>0)<br>• in minimum two independent experiments | • ≥11 / ≤ 71 codons<br>• peak height ≥ 20<br>• no Ribo peak or log2FC ≥1<br>• in minimum two replicates<br>• add top 16 candidates 4-10 codons | • shortest ORF ≤ 71 codons<br>• longest ORF ≥ 11 codons<br>• no TIS(Onc) peak allowed<br>• excl. internal out of frame and within 25 nt downstream of annotated stop codon |
| | **manual curation** | | |

**Supplementary Figure S8. Establishing cutoffs for novel CJsORF predictions from Ribo-seq data. (A)** Transcript levels (RPKM, *left*), translational efficiency (TE, *middle*), and DeepRibo score (*right*) for the validated (based on MS and tagging and WB analysis) sORF benchmark set (blue) vs. not validated sORFs (black). Dashed lines: cutoffs used for predictions. Solid lines: Mean within each sORF set. Ribo-seq(Cm) is shown as a representative experiment. For DeepRibo scores, higher values indicate higher confidence predictions (Clauwaert et al., 2019). sORFs without predictions are indicated below (light blue/grey dots, "not detected"). **(B)** TIS peak height (reads, *left*) and corresponding TIS/Ribo peak ratio (log2FC, *right*) for validated sORF benchmark set (blue) and not validated sORFs (black). Dashed lines: cutoffs for sORF predictions. Solid lines: Mean within each sORF set. TIS(Ret) was used as a representative experiment. sORFs without a TIS peak (left graph) are indicated below (light blue/grey dots). Those without a Ribo peak (TIS peak only, "no log2FC", right graph) are shown above (blue/grey dots). sORFs without a TIS peak were not considered in the right plot. **(C)** Comparison of TTS predictions from TTS(Mono) and TTS(Di) data for the sORF benchmark set. **(D)** Overview of filtering criteria for sORF candidates from all datasets. Related to main **Fig. 5A**.

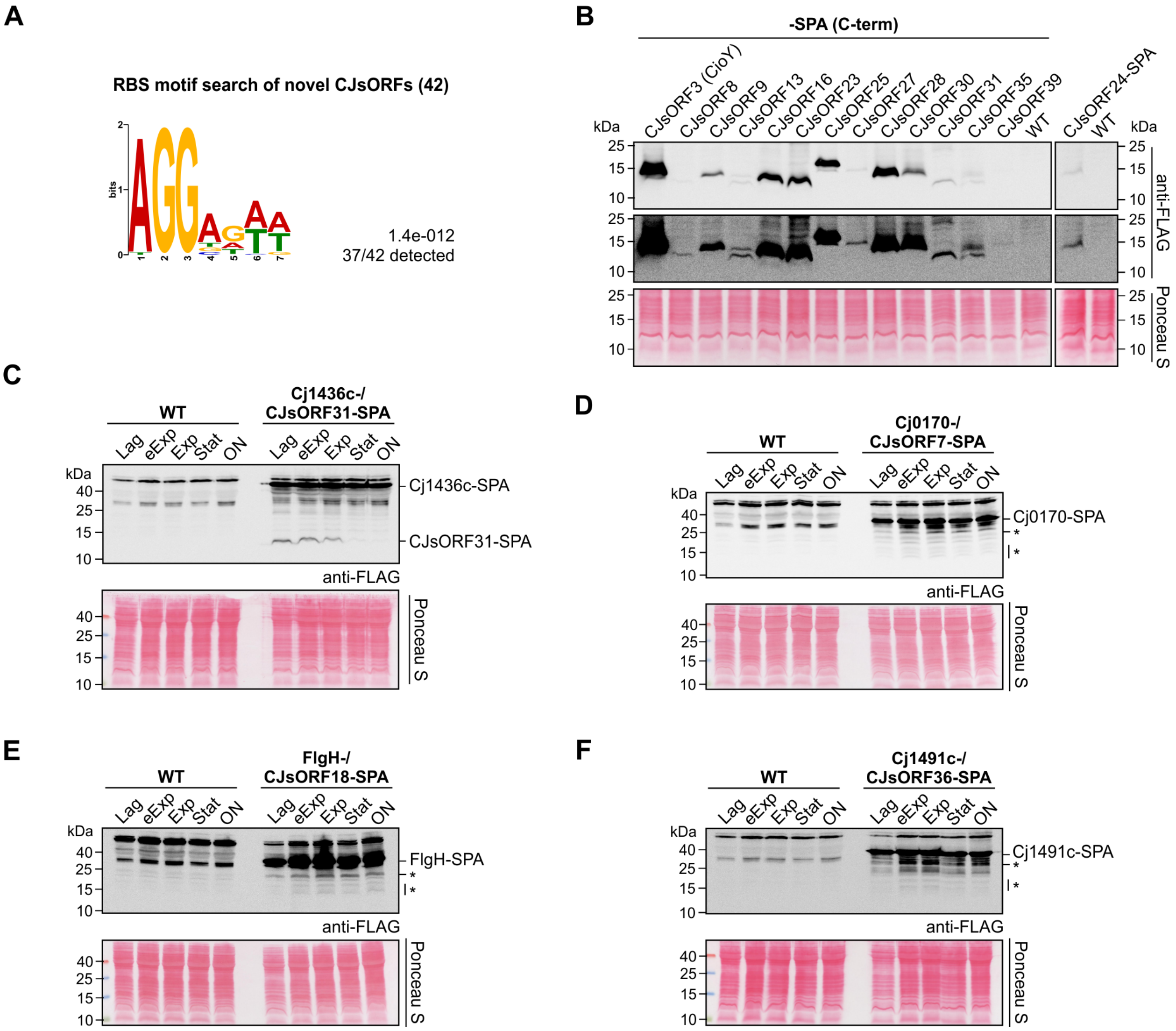


**Supplementary Figure S9. Additional validation of novel CJsORFs. (A)** MEME analysis of 15 nt upstream of novel CJsORFs (https://meme-suite.org/meme/ (Bailey et al., 2009)). **(B)** Parallel WB analysis of all detected novel CJsORFs. Samples were harvested from lag phase (at $OD_{600}$ 0.1) from strains carrying a C-terminal SPA tag at the native locus. Middle panel: higher exposure of top panel. **(C)** Full WB for CJsORF31 (related to main **Fig. 5B**). **(D-F)** Full WB for internal in-frame candidates CJsORF7, CJsORF18, and CJsORF36. Parental genes are indicated, as introduction of the epitope at the sORF C-terminus also tags the parental gene. Asterisk: potential CJsORF bands. eExp: early exponential. Stat: Stationary. ON: overnight. For WB, SPA-tagged proteins were detected from 0.2 $OD_{600}$ equivalents of cell lysate with an anti-FLAG antibody. WT: untagged WT antibody negative control. Ponceau S staining of membranes served as a loading control. WB are representative of at least two independent experiments.

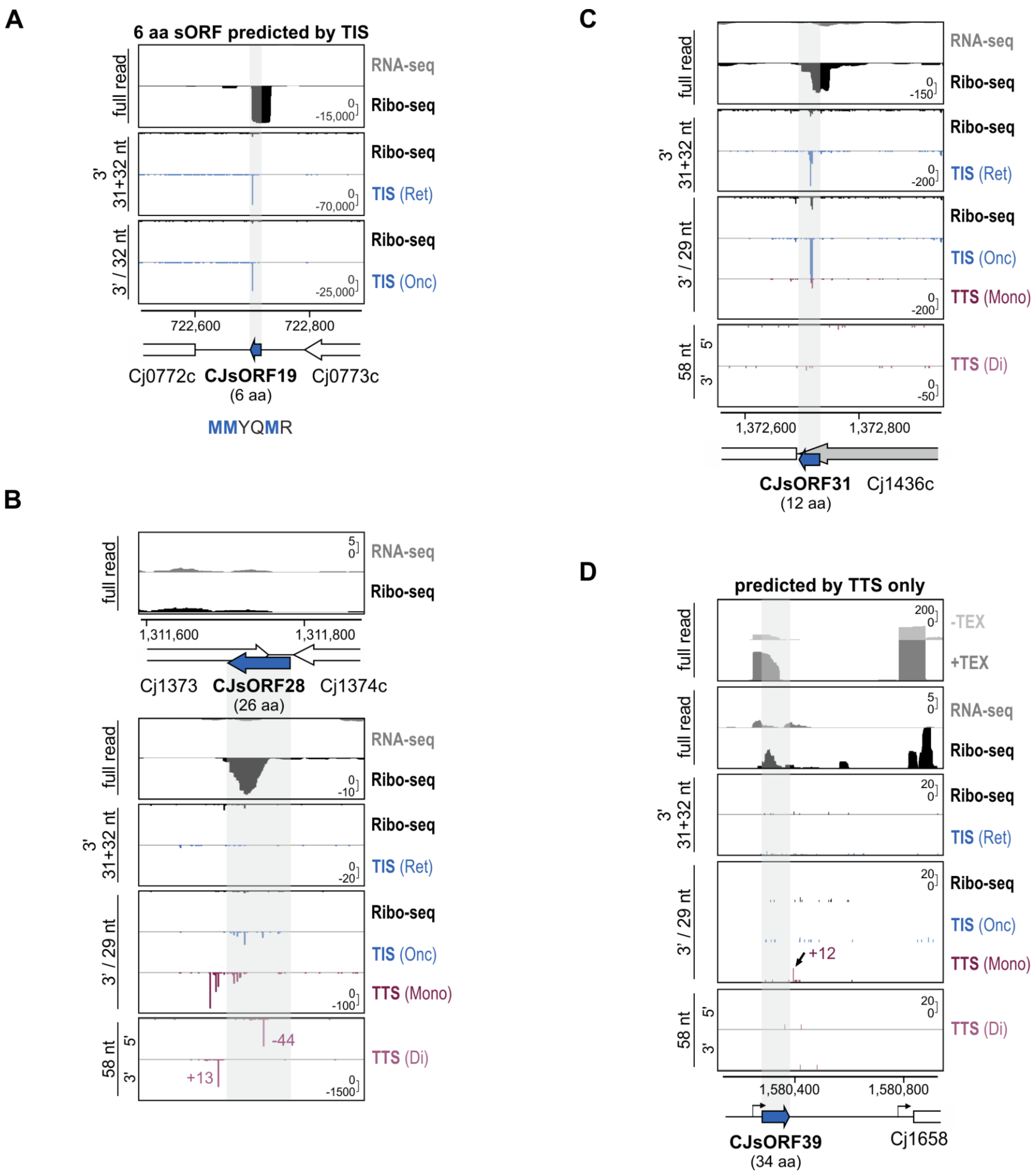


**Supplementary Figure S10. Novel CJsORFs predicted from TIS (Ret and Onc) and TTS profiling data. (A)** A short ORF (CJsORF19, 6 aa) was among the top TIS candidates between 4 and 10 codons in length encoded next to genes involved in methionine transport. The CJsORF19 is enriched for methionine codons. **(B & C)** Full coverage for novel sORFs CJsORF28 and CJsORF31. Related to main **Fig. 5E**. **(D)** The intergenic novel sORF CJsORF39 was derived from TTS predictions alone. Bent arrows: TSS based on dRNA-seq (Dugar et al., 2013). Y-axis: rpm (reads per million).

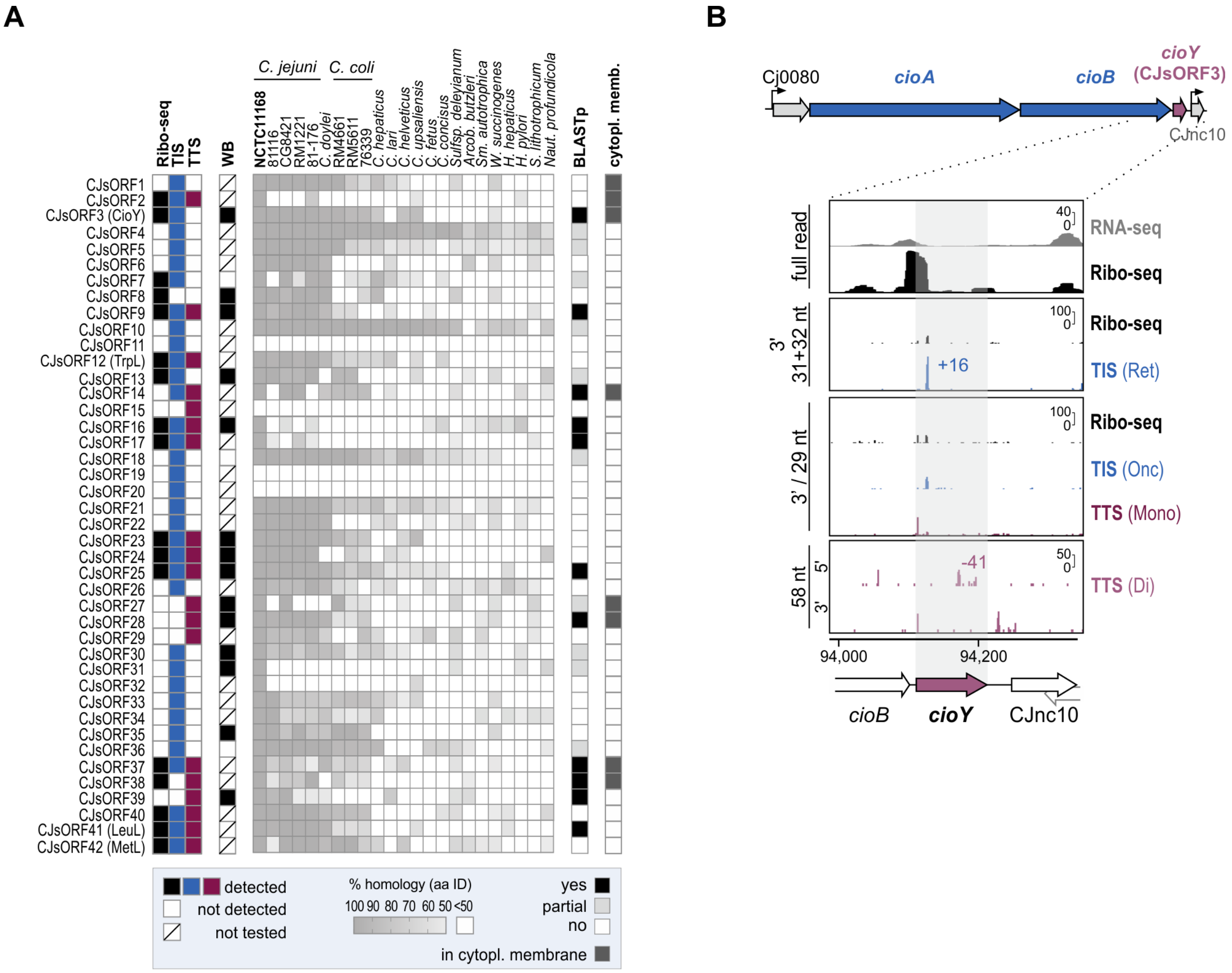


**Supplementary Figure S11. Conservation of novel sORF candidates in Epsilonproteobacteria reveals a terminal oxidase small protein component *cioY*. (A)** Extended conservation analysis of novel *C. jejuni* small proteins. Related to main **Fig. 6A**. WB: Epitope-tagged version detected by western blot at any growth phase. MS: detected by mass spectrometry in log phase. Cytoplasmic membrane (Cytopl. memb.) localisation was predicted by pSORTb (Yu et al., 2010). Homology (percent amino acid identity, aa ID) was determined by tBLASTn searches at NCBI using the *C. jejuni* NCTC11168 sequence as query and parameters as described in the Methods. BLASTp: yes - at least one 100% match in length and sequence. partial - sequence aligns with part of an (often longer) annotated ORF (see also **Table S5**). **(B)** Translatomics coverage for *cioY* (CJsORF3, 34 aa) located in the conserved Cj0080-*cioA*-*cioB*-*cioY* operon. Bent arrows: TSS based on dRNA-seq (Dugar et al., 2013). Y-axis: rpm (reads per million).

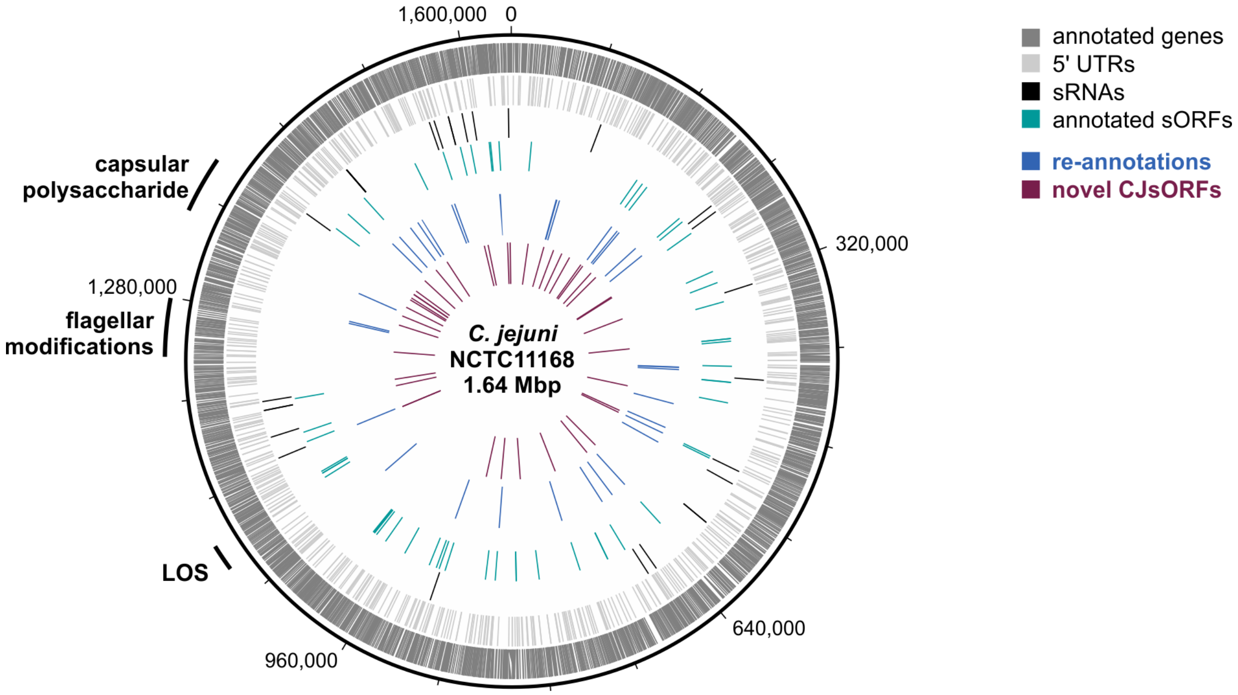


**Supplementary Figure S12: Overview of the refined annotation for the *C. jejuni* NCTC11168 chromosome (NC_002163.1).** Genome ring displaying new or updated annotated features as well as previously published 5'UTR and sRNA annotations (Dugar et al., 2013). Dark grey: NCBI annotation. Light grey/black: 5′UTRs/sRNAs (Dugar et al., 2013). Green: sORFs (≤ 70 aa) in NCBI annotation. Blue: re-annotations/additions based on this study. Red: novel CJsORFs identified in this study. Hypervariable LOS (lipooligosaccharide) biosynthesis, capsular polysaccharide biosynthesis, and flagellar modification gene clusters are indicated (Parkhill et al., 2000). Genomic features were visualized with Artemis DNA Plotter (Carver et al., 2009).

### Supplementary Methods

#### Transformation of *C. jejuni* mutant construction

Strains grown from frozen stocks until passage one or two on MH agar were harvested into cold electroporation solution (272 mM sucrose, 15% (v/v) glycerol) and washed twice with the same buffer. Cells (50 µl) were mixed with 500-1000 ng PCR product on ice and electroporated (Bio-rad MicroPulser) in a 1 mm gap cuvette at 2.5 kV. Cells were then transferred with *Brucella* broth to a non-selective MH plate and recovered overnight at 37°C micro-aerobically before plating on the appropriate selective medium. Strains were validated by colony PCR and for complementation, overexpression, and epitope tagging, validated by Sanger sequencing (Macrogen, Microsynth).

***C. jejuni* non-polar deletion mutant construction by recombination with overlap PCR products.** Non-polar deletion mutants of protein coding-genes were constructed by homologous recombination with overlap PCR products consisting of a resistance cassette in between approximately 500 bp of sequence upstream and downstream of the target gene using primers listed in **Table S9**. As an example, deletion of *cmeB* with a non-polar $Gm^R$ cassette is described. An approximately 500 bp region upstream of *cmeB* (Cj0366c) was amplified using CSO-4091/4092, while the downstream region was amplified using CSO-4093/4094. The 5′ ends of the antisense primer for the upstream region and the sense primer for the downstream region included regions overlapping the resistance cassette 5′ and 3′ end, respectively. The $Gm^R$ cassette was amplified using primers HPK1/HPK2 from pUC1813-apra (Bury-Moné et al., 2003). Next, a three-fragment overlap PCR was performed using the *cmeB* upstream, *cmeB* downstream, and $Gm^R$ cassette fragments in an equimolar ratio and primers CSO-4091/4094. Following confirmation of the correct size by gel electrophoresis, the resulting overlap PCR product was electroporated as described above into WT and deletion mutants were selected on plates containing gentamicin. The deletion strain (Δ*cmeB*, CSS-5613) was confirmed using a primer binding upstream of the *cmeB* upstream fragment (CSO-4091) and an antisense primer binding the $Gm^R$ cassette (HPK2). For non-polar $Gm^R$, $Kan^R$, and $Hyg^R$ deletions, cassettes were amplified using

HPK1/HPK2 from pUC1813-apra (Bury-Moné et al., 2003), pGG1 (Dugar et al., 2016), or pACH1 (Cameron and Gaynor, 2014) as template, respectively. For PCR reactions including the $Hyg^R$ cassette, 3% DMSO was included in reactions.

**C-terminal 3×FLAG-, SPA-, or sfGFP-tagging.** Epitope tagging was performed via homologous recombination at the native locus. The overlap PCR product contained approximately 500 bp upstream of the sORF penultimate codon fused to the 3×FLAG, SPA, or sfGFP sequence, a $Kan^R$ or $Gm^R$ resistance cassette, and the ORF downstream region. As an example, tagging of Cj0978c with 3×FLAG is provided. The Cj0978c upstream and coding region was amplified using CSO-3344/3770, and 500 bp of the downstream was amplified with CSO-3346/3347. A fusion of 3×FLAG to the $Gm^R$ cassette was amplified using CSO-0065/HPK2 on the previously published 3×FLAG strain (*fliW*::3×FLAG) (Dugar et al., 2016). The up, down, and 3×FLAG-$Kan^R$ fragments were mixed together in approximately equimolar ratios, annealed, and amplified by overlap PCR using primers CSO-3344/3347. Clones were validated by colony PCR with CSO-3348/HPK2 and sequencing with CSO-0023 or CSO-3348.

**Heterologous expression from *rdxA*.** The *rdxA* locus (Cj1066) can be used for heterologous gene expression in *C. jejuni* (Ribardo et al., 2010). Constructs for complementation in *rdxA* were made mostly in plasmids containing approximately 500 bp of upstream and downstream sequence from *rdxA* flanking a $Cm^R$ or $Kan^R$ cassette (with promoter and terminator) by subcloning the *C. jejuni* sequence into previously-constructed plasmid vectors based on pFK8.5, pSE59.1, or pGD34.7 (Alzheimer et al., 2020; Dugar et al., 2018). To generate pMA16.1 (for heterologous expression of *pgp1* from *rdxA*) the backbone of pGD34.7 was amplified with CSO-3641/0350 and two inserts were amplified from 81-176 WT (CSS-0063) gDNA with CSO-3646/3647 (promoter region) and Cjj81176_1344 (*pgp1*-A8) with CSO-3644/3645. Both insert fragments were fused via overlap PCR with CSO-3646/3645, digested with *Nde*I/*Xma*I, and ligated into the plasmid backbone. Positive clones were identified by colony PCR with CSO-0463/3270 and sequencing with CSO-0646/3270. For Cj0455c complementation, inserts for Cj0455c full-length were amplified with CSO-5851/5852 or for Cj0455c truncated with CSO-5851/5853 from WT gDNA (CSS-

5295), digested with *Bam*HI and ligated into a backbone amplified from pSE59.1 with CSO-5284/1354). Positive clones were identified by sequencing with CSO-3270. Alternatively, overlap PCR was used for, *e.g.*, CJsORF35-SPA. The backbone for generation of pFK8.5 was amplified from pST7.2 (similar to published pST1.1 (Dugar et al., 2018)) with CSO-0347/4738 and digested with *Cla*I. The insert (CJnc230 sRNA, (Dugar et al., 2013)) was amplified with CSO-4254/4257 from WT gDNA (CSS-5295) and also digested with *Cla*I. Vector and insert were ligated, transformed into *E. coli* TOP10 cells and validated by colony PCR with CSO-2276/4257. Next, a *rdxA*_UP-Kan$^R$-P$_{porA}$ was amplified with CSO-2276/4738, the *rdxA*_DN region with CSO-1785/2277 from pFK8.5, and the SPA sequence was amplified with CSNIH-0080/CSO-4891 from pJL148. The CJsORF35 insert (including 103 nt upstream) was amplified from WT gDNA (CSS-5295) with CSO-4889/4890. The four fragments were mixed in an approximately equimolar ratio and subjected to overlap-extension PCR with CSO-2276/2277. This product was used for electroporation. Primers CSO-2276/2277 were used to amplify all constructs from *rdxA*-based plasmids for electroporation into *C. jejuni*. Clones with intended insertions were validated by colony PCR using CSO-0643/0644/0349 (*rdxA*-Cm$^R$ constructs) or CSO-0023/0349 (*rdxA*-Kan$^R$ constructs). Insertions were sequenced with CSO-3270, CSO-0643, CSO-0644, or CSO-0023.

**Growth and harvest for ribosome profiling (Cm).** Translation was arrested in WT *C. jejuni* cultures and cells harvested for Ribo-seq using Cm ice as described previously with modifications (Becker et al., 2013; Oh et al., 2011). *C. jejuni* NCTC11168 WT cultures were grown to mid-log phase ($OD_{600}$ approx. 0.5) in 100 ml BB medium at 37˚C with shaking at 150 rpm under microaerobic conditions. A sample for total RNA was transferred to RNA stop mix (95% ethanol, 5% buffer-saturated phenol (Roth)) and snap-frozen in liquid $N_2$. Bacteria were then treated with 1 mg/ml Cm (Sigma) for 5 min at 37°C under microaerobic conditions, followed by immediate chilling by mixture with an equal volume of crushed ice containing 1 mg/ml Cm and incubation on ice for 10 minutes. Cells were harvested by centrifugation at 10,000 *g* for 10 min, and immediately frozen in liquid $N_2$.

**Minimum inhibitory concentration (MIC) determination for TIS/TTS profiling antibiotics.** To determine the sensitivity of *C. jejuni* strains to antibiotics used for TIS/TTS profiling, we measured their MIC (Wiegand et al., 2008) in the same broth as used for Ribo-seq using a 96 well-plate format with serial 2-fold dilutions of antibiotic. Briefly, strains from an overnight culture in BB with vancomycin (WT (CSS-5295), Δ*cmeB* (CSS-5617) or ΔCj0182 (CSS-4077, *sbmA* homolog)) were diluted into fresh medium to an $OD_{600}$ of 0.0005. Antibiotics (Ret - Sigma CDS023386, Onc & Api - NovoPro BioScience Inc., Shanghai, China) were diluted serially by 2-fold in BB with vancomycin, starting at a final concentration (including bacterial suspension) of 8 µg/ml for tetracycline (Carl Roth Art. No. 0237.2) or Ret, or 16 µM PrAMPs. After addition of bacteria to the wells, plates were incubated under microaerobic conditions at 37°C for 24 hours. The MIC was taken as the lowest concentration where no visible bacterial growth was observed.

| | **WT** | **Δ*cmeB*** | **ΔCj0182** |
|---|---|---|---|
| Tetracycline (µg/ml) | 0.0625 | 0.031 | 0.031 |
| Retapamulin (µg/ml) | 0.125 | 0.016 | N/A |
| Oncocin (µM) | 1.56-0.78 | N/A | 50 |
| Apidaecin-137 (µM) | 6.25 | N/A | >50 |

**Growth and harvest for TIS profiling (Ret).** *C. jejuni* Δ*cmeB* was grown, treated, and harvested for TIS profiling as described for Cm Ribo-seq above with minor modifications based on protocols established previously in *E. coli* (Meydan et al., 2019; Weaver et al., 2019). Two cultures were grown in parallel until exponential growth phase in BB medium, and a sample was removed, mixed with RNA stop mix, and immediately frozen in liquid $N_2$ for total RNA analysis. One culture was then treated with Ret (12.5 µg/ml final concentration) for 10 minutes under routine growth conditions. Cells were then harvested by fast-filtration with a 0.45 µm polyethersulfone membrane (Millipore) and immediately frozen in liquid $N_2$.

**Growth and harvest for TIS profiling (Onc) and TTS.** For the TIS/TTS(Onc/Api) experiment, *C. jejuni* WT was grown in BB medium until log phase ($OD_{600}$ 0.4) in three parallel cultures. A sample was removed for total RNA analysis, mixed with RNA stop mix, and immediately frozen in liquid $N_2$. One culture was left untreated, while one was treated with Onc or Api (50 µM final concentration) for 10 min under routine growth conditions, respectively. Cultures were then immediately transferred to pre-chilled glass flasks and swirled in an ice bath for 3 min to rapidly chill the cells and halt translation. Chilled cells were harvested by centrifugation at 4500 rpm and immediately frozen in liquid $N_2$.

**Detection of TIS/TTS sites and associated ORFs.** For detection of TIS/TTS, we adapted previous peak detection methods (Gelsinger et al., 2020; Weaver et al., 2019). All programming scripts are available at https://github.com/RickGelhausen/StartStopFinder. The lack of support for TTS libraries at the time provided the impetus to develop our own tool rather than previously published scripts (Bartholomäus et al., 2021). Our method also avoids issues with overlapping start codons that were not addressed in these scripts. First, coverage files (normalized per million reads) were generated by HRIBO (Gelhausen et al., 2021) using different single-nucleotide mapping strategies (3′ or 5′ end) and read lengths (27-33 nt). Metagene analysis of ribosome occupancy at start or stop codons was performed as described previously (Becker et al., 2013). Read length and offsets (see below) were determined independently for each experiment (*i.e*, TIS(Ret) and TIS/Onc).

These parameters were used for TIS peak detection and ORF prediction. All start codons (ATG/TTG/GTG) were collected and intervals of 5 nt, selected based on metagene analysis, that span around each start codon were generated. Intervals were then shifted by the previously determined offset. For each position in the coverage file that overlapped a given interval, the peak height was defined as the sum of the overlapping positions that had a read count higher than 5 reads. For each interval that had a non-zero peak height, the next in-frame stop codon was identified to generate a corresponding ORF (of any length). Associated TE and RPKM values were calculated. A similar method was used for detection of stop codons with TTS data, except potential sORFs with stop codons downstream of annotated genes within 25 nt were excluded to reduce signals from ribosome stop codon readthrough.

The following coverage files and offsets were used for start codon detection: TIS(Ret): 3′ end read coverage, +16 nt offset, 31 + 32 nt read length coverage files; TIS/TTS(Onc/Api): 3′ end of read coverage, +17 nt offset, 32 nt read length coverage files. For peak detection in TTS libraries using the following data: TTS-Mono: 3′ end of read coverage, +13 nt offset, 29 nt read length coverage files; TTS(Di): 3′ end of read coverage, +13 nt offset, 58 nt read length coverage files or 5′ end of read coverage, -45 nt offset, 58 nt read length coverage files. For start codons, offset positions represent the 16th nt after the first nt of NTG (*e.g*., for + 16 nt). For stop codons, offset positions represent the 13th nt after the first nt of the stop codon (*e.g*., for + 13 nt). Predictions were run with similar parameters as for TIS detection, except an interval of ± 3 nt around the offset nucleotide was used.

TIS and TTS sites were classified as follows: Annotated: within 3 nt (up or downstream) of an annotated Start codon; Internal in-frame: within an annotated ORF, N-terminal truncation/Internal start site and same stop codon; Internal out-of-frame: within an annotated ORF, different reading frame; Unannotated - outside of annotated features.

sORF predictions based on the third experiment (TIS(Onc) and TTS) were generated using only a single replicate available at the time. To validate the method and determine the robustness of TTS detection, we generated two additional replicates and used these for automated predictions of annotated and novel sORFs (**Table S10**, predictions summarized in **Table S11**). This data was also used to manually curate TTS for longer genes (*e.g*., phase-variable, **Fig. 4 & Table S11**).

**Mass spectrometry based proteomics.** For preparation of soluble *C. jejuni* protein extracts, cells were disrupted in a FastPrep Homogenizer (MP-Biomedicals) for 30 s at 6.5 m/s$^2$ followed by incubation on ice for 4 min. This procedure was repeated two times. Immediately after the third disruption step, the lysate was centrifuged two times for 15 min (12,000 g at 4°C) to remove all cell debris and insoluble and aggregated proteins. Protein concentration was determined as described previously (Fuchs et al., 2021). Two different techniques for pre-fractionation of proteins were applied: (i) separation of soluble proteins by one dimensional (1D) SDS-PAGE and in-gel digestion with trypsin or chymotrypsin (see "gel-based approach" described in (Fuchs et al. 2021), or (ii) fractionation of proteins on a GELFREE 8100 fractionator (Expedeon) and trypsin digestion. For the gel-based approach, we separated 36 µg protein crude extracts on 15% polyacrylamide gels. For in-gel digestion with chymotrypsin, an enzyme/protein ratio of 1:10 was applied for 16 h at pH 8 and 30°C.

For protein fractionation on a GELFREE 8100 fractionator, 200 µg soluble proteins were separated according to manufacturer's instructions (Expedeon) using 10% Tris acetate cartridges. In addition to the samples recommended by the manufacturer (Expedeon), a sample 220 min after starting fractionation was taken. Protein digestion was performed in protein low binding tubes (Eppendorf, Hamburg, Germany) using the Single-Pot Solid-Phase-enhanced Sample Preparation technique described previously (Hughes et al., 2014) with some modifications. In brief, magnetics beads were washed three times with MilliQ water before use. 30 µg beads were added to each sample, which was adjusted to pH <5 using 5% formic acid. To bind the proteins to the beads, a fourfold volume of acetonitrile was added and samples were mixed horizontally at room temperature. After two to three hours, additional 30 µg beads were given to each sample which were incubated as aforementioned overnight. Samples were centrifuged (13,000 g, 5 min, room temperature), placed on a magnetic rack and the supernatant was removed. Proteins bound to magnetic beads were washed two times with ethanol and incubated with 50 mM DTT in 50 mM $NH_4HCO_3$, 1 mM $CaCl_2$ for 30 min at 60°C. Subsequently, iodoacetamide was added to each sample with a final concentration of 120 mM and incubated for 20 min at room temperature. For protein digestion, trypsin was applied with an enzyme/protein ratio of 1:50. After incubation for 12 h (800 rpm, 37 °C), samples were adjusted to pH <5 using 5% formic acid, centrifuged and peptides bound to magnetic beads were washed two times with acetonitrile. Peptide elution was performed in two steps. First, magnetic beads were treated with 20 µl 2% DMSO for 30 min, centrifuged and the supernatant was transferred to a new tube. In a second step, the beads were incubated with 20 µl 0.065% formic acid, 500 mM KCl in 30% acetonitrile for 30 min and centrifuged. The supernatants of both elution steps were combined, vacuum dried and stored at -20°C. Peptide desalting was performed as described previously (Fuchs et al., 2021).

Peptide fractions were analyzed using the Orbitrap Fusion MS coupled to a Dionex Ultimate 3000 nHPLC system (Thermo Fisher Scientific Inc., Waltham, Massachusetts, USA) as described by Fuchs *et al.* (Fuchs et al., 2021) with some modifications. Primary Scans were performed in the profile modus scanning an m/z of 350 - 1,700 with a resolution (full width at half maximum at m/z 400) of 120,000 and a lock mass 445.12003. Using Xcalibur software (Thermo Fisher Scientific Inc., San Jose, SA, USA), the mass spectrometer was controlled and operated in the "top 20" mode, selecting the 20 most abundant MS ions for fragmentation. Primary ions (±10 ppm) were selected by the quadrupole (isolation window:

1.6 m/z), fragmented by CID (collision energy 35%, activation Q 0.25) and analyzed in the ion trap with an exclusion time of 20 s. The charges of MS ions used for fragmentation were 2 to 6 for trypsin or 1 to 6 for chymotrypsin.

For identification of small proteins in *C. jejuni* based on MS/MS data, we used the bacterial proteogenomics workflow described previously (SALT & Pepper; https://gitlab.com/s.fuchs/pepper; (Fuchs et al., 2021)). This fully automated workflow includes protein database generation, database searching, peptide-to-genome mapping, and result interpretation. MS- and MS/MS-data of all samples were searched by MaxQuant (Max Planck Institute of Biochemistry, Martinsried, Germany, www.maxquant.org, version 1.5.2.8) against a database with *C. jejuni* annotated protein sequences from NCBI (downloaded at 01-09-2020) and sORFs that were predicted using our Ribo-seq data and a translational database (TRDB) of the full coding potential of the *C. jejuni* genome generated by six-frame translation from stop codon to stop codon with a minimum length of 9 aa generated by SALT (https://gitlab.com/s.fuchs/pepper). For chymotrypsin, the number of missed cleavages was set to 4 and maximum charge to 7.

**Co-immunoprecipitation (coIP) for investigation of protein-protein interactions.** Strains carrying chromosomally epitope-tagged versions of CioA and/or CioY were used (CioA-SPA & CioY-sfGFP (CSS-8004) and reciprocal version CioY-SPA & CioA-sfGFP (CSS-6727)). Pulldown of the SPA-tagged protein was performed with an anti-FLAG antibody (Sigma-Aldrich, #F1804-1MG) bound to Protein A-Sepharose beads (Sigma-Aldrich, #P6649), and co-purification of the second sfGFP tagged protein was investigated by western blot with an anti-GFP antibody (Roche #11814460001). As a control for unspecific binding, lysates of an untagged wild-type strain (CSS-5295) as well as the corresponding sfGFP-only tagged strain (CioA-sfGFP (CSS-6725) or CioY-sfGFP (CSS-8002)) alone were also used. Lysates for coIP were prepared from ~60 $OD_{600}$ of cells harvested at exponential phase ($OD_{600}$ 0.6, 5000 rpm, 20 min, 4°C) which were first washed once in buffer A (20 mM TrisHCl pH 8.0, 1 mM $MgCl_2$, 150 mM KCl, 1 mM DTT) (8000 *g*, 2 min, 4°C). In parallel, 1 $OD_{600}$ of cells was harvested for whole cell lysate analysis and boiled for 8 min at 95°C in 1× protein loading buffer. After the supernatant was discarded, cell pellets were snap frozen in liquid nitrogen and stored at -80°C until further use. Thawed cell pellets were lysed in 1 ml lysis buffer (buffer A incl. 1 mM PMSF (phenylmethylsulfonyl fluoride, Roche), 20 U DNase I (Thermo Fisher Scientific), 200 U RNase Inhibitor (moloX, Berlin) and 1% DDM (n-dodecyl-B-D-maltoside) with a FastPrep system (MP Biomedical, matrix B, 1× 4 m/s, 10 sec). Lysates were cleared by centrifugation (13,000 rpm, 10 min, 4°C) and an aliquot (1 $OD_{600}$) was set aside as the input/lysate control for western blot analysis. The lysate was then incubated with rotation for 30 min at 4°C with 35 µl anti-FLAG antibody. The Protein A-Sepharose beads (75 µl/sample) were washed three times with buffer A and then lysates, pre-incubated with anti-FLAG antibody, were added to the pre-washed beads and incubated for another 30 min at 4°C with rotation. The supernatant (unbound fraction) was removed after centrifugation (15,000 *g*, 1 min, 4°C). Beads with the bound proteins were washed five times with buffer A. Finally, the bound proteins were eluted by adding 400 µL 1× protein loading buffer and boiling for 8 min at 95°C with shaking at 1000 rpm. The eluate was precipitated with a minimum 6 vol acetone overnight at -20°C. Precipitated proteins were collected by centrifugation (15,000 rpm, 1 h, 4°C), dried, and resuspended in 1× protein loading buffer. For verification of successful coIP and to investigate potential interactions, whole cell lysate (0.1 $OD_{600}$), lysate/input samples (0.1 $OD_{600}$) and the eluates (5 $OD_{600}$) were used for western blot analysis. Blots were probed for anti-GFP for investigation of interaction, anti-FLAG for verification of pull-down, and finally anti-GroEL as a loading control. Two independent biological replicates of each coIP experiment were performed.
